# Discovery of potent oligopeptides for various metabolic diseases using deep learning

**DOI:** 10.64898/2026.01.06.697667

**Authors:** Baichuan Xiao, Hao Zhu, Chao Ma, Xiaoran Wang, Zhanying Feng, Ruirui Lang, Hailin Shan, Ziqiu Meng, Runbang He, Yixiang Zhou, Jian Huang, Siyang Wei, Daiyi Liu, Jiqiang Liu, Zhenni Shi, Xiaokai Tang, Jinmo Wang, Chao Liu, Liguo Wang, Zhanzhan Li, Yong-Biao Zhang

**Affiliations:** School of Engineering Medicine, Beihang University, Beijing, 100191, China; The First College of Clinical Medicine, Henan University of Chinese Medicine, Zhengzhou, 450046, China; Department of Statistics, Department of Biomedical Data Science, Stanford University, Stanford, CA 94305, USA; Tianjin Institutes of Health Science, Tianjin, 300000, China; Division of Computational Biology, Mayo Clinic College of Medicine and Science, Rochester, MN 55905, USA; Academy of Traditional Chinese Medicine, Henan University of Chinese Medicine, Zhengzhou, 450046, China; Key Laboratory of Big Data-Based Precision Medicine and Key Laboratory of Innovation and Transformation of Advanced Medical Devices, Ministry of Industry and Information Technology; Key Laboratory of Biomechanics and Mechanobiology, Ministry of Education, Beijing, China; National Medical Innovation Platform for Industry-Education Integration in Advanced Medical Devices (Interdiscipline of Medicine and Engineering), Beihang University, Beijing, 100083, China

**Author notes:** Correspondence Yong-Biao Zhang, Address: 37 Xueyuan Road, Haidian District, Beijing, P.R. China, 100191, Zhanzhan Li, Address: 156 Jinshuidong Road, zhengzhoudong District, Zhengzhou, P.R. China, 450046. These authors contributed equally to this work (Baichuan Xiao, Hao Zhu, Chao Ma, Xiaoran Wang).

## Abstract

Artificial intelligence (AI)-based methods are increasingly critical in peptide drug discovery, but their applications are limited in a narrow scope such as antimicrobial peptides. For indications where peptide therapy inherently excels, such as metabolism, endocrinology, and tissue regeneration, AI-based pipeline for therapeutic peptide discovery is long awaited but still unmet. Here, we propose a deep learning-based pipeline, Deepeptide, capable of identifying therapeutic oligopeptides for various metabolism-related indications. Leveraging the intrinsical relationship of ‘disease indication — biological processes — molecular functions’, Deepeptide discovers oligopeptides with indication-ameliorating-related molecular functions as lead candidates for indication of interest. Deepeptide was applied in five representative indications of metabolism, endocrinology, and tissue regeneration: angiogenesis, lipid metabolism, osteogenesis, glucose metabolism, and anti-angiogenesis. Overall, 62% of the identified oligopeptide candidates demonstrated significant bioactivity in vitro, with most of them showing comparable potency to the first-line drugs. Notably, the heptapeptide AP7 exhibited angiogenic potency comparable to VEGF in excisional wound splinting mouse model by promoting cell migration rather than proliferation, and hexapeptide TP6 showed significant dual-efficacy against hyperlipidemia and obesity in high-fat diet mice by inhibiting lipid synthesis and regulating gut microbiota. These findings highlight the potential and generalizability of Deepeptide in therapeutic oligopeptide discovery for metabolic diseases.

## INTRODUCTION

Peptide drugs have drawn significant attention from drug developers due to their advantages of high potency, selectivity, and generally low toxicity and immunogenicity^1–3^. As key biological mediators, peptides have shown unique strengths in treating a range of indications, such as endocrinology, metabolism, and oncology^3^. Among these, oligopeptide drugs, as a major subgroup, offer additional benefits, including ease of synthesis and cost-effectiveness^2–4^. However, discovering therapeutic oligopeptides for the majority of disease indications, even those where peptides have demonstrated unique advantages, remains a significant challenge^1,4,5^.

In recent years, AI-driven drug discovery has made remarkable strides across various fields, including peptide drugs^1,6,7^. However, these efforts have primarily focused on indications with sufficient training data, such as antimicrobial peptide^8–10^, anticancer peptide^11–13^, and cell-penetrating peptide^14–16^. Moreover, the pace of positive data acquisition and accumulation has been relatively slow, resulting in a huge application gap for most indications that lack sufficient positive training data, especially for these metabolism-related disease indications^5,17^. Therefore, an AI pipeline for therapeutic oligopeptide discovery applicable for various metabolic diseases is long awaited but still unmet.

Here, we developed Deepeptide, an AI-based generalizable pipeline capable of discovering potent and novel oligopeptides for various metabolic diseases. Inspired by the intrinsical relationship of ‘disease indication — biological processes — molecular functions’, Deepeptide discovers oligopeptides with indication-ameliorating-related molecular functions as lead candidates for indication of interest, circumventing the requirement for indication-specific training data. Specifically, Deepeptide implements its generalizable pipeline by two-steps. First, it constructs a specialized library composed of protopeptides with indication-ameliorating-related molecular functions, in which oligopeptide candidates are contained as sub-fragments. Second, it extracts oligopeptide candidates from these protopeptides using a deep learning (DL) model and prioritizes these candidates by inferring their indication-ameliorating-related molecular functions. The top-prioritized ones are recommended as therapeutic oligopeptide candidates for subsequent experiment validations.

Deepeptide was applied in five representative disease indications of metabolism, endocrinology, and tissue regeneration. Overall, 62% of these candidates demonstrated significant bioactivity in vitro, and most of them showed comparable potency to first-line drugs. Notably, heptapeptide AP7 exhibited angiogenic potency comparable to VEGF in an excisional wound splinting mouse model by promoting cell migration rather than proliferation, and hexapeptide TP6 showed significant dual-efficacy against hyperlipidemia and obesity in high-fat diet mice by inhibiting lipid synthesis and regulating gut microbiota. These findings highlight the capacity and generalizability of Deepeptide in therapeutic oligopeptide lead discovery for metabolic diseases.

## DESIGN

The fundamental design principle of Deepeptide is: disease indications are intrinsically linked to the biological processes of proteins, which are underpinned by molecular functions that are often executed by specific fragments of protein like oligopeptides^18,19^. This knowledge provides a feasible guideline for identifying oligopeptides with indication-ameliorating-related molecular functions as therapeutic oligopeptides for indications of interest (**Figure 1A**).

**Figure 1.**
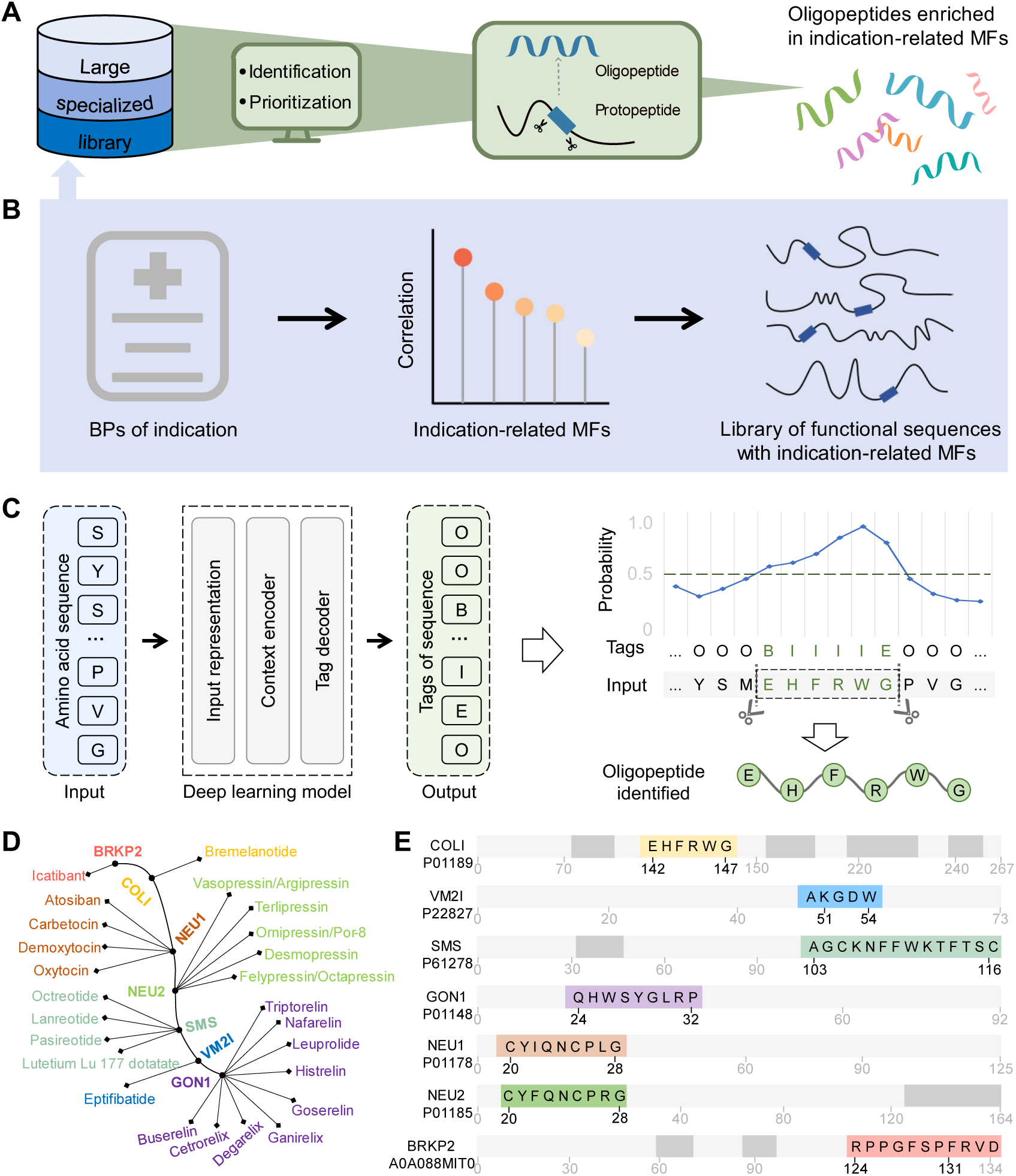
Overview of the Deepeptide pipeline. (**A**) A flowchart depicts the two-step pipeline of Deepeptide. Deepeptide identifies oligopeptides with indication-ameliorating-related molecular functions as lead candidates for indication of interest, by first constructing a large specialized library of functional protopeptides (blue section) and then identifying and prioritizing oligopeptides from the library (green section). (**B**) A schematic diagram illustrates the construction of a specialized library composed of protopeptides with indication-ameliorating-related molecular functions, based on the intrinsical relationship among disease indications, biological processes, and molecular functions. (**C**) Left: The architecture of the deep learning (DL) model designed to identify biopeptide candidates from input amino acid sequences. Right: Tags of sequence are used to extract oligopeptides from protopeptides. Tags ‘B’, ‘I’, and ‘E’ denote the beginning, inner part, and end of an oligopeptide candidate, respectively, while ‘O’ indicates regions outside of the oligopeptide candidate. (**D**) Twenty-five marketed oligopeptide drugs are grouped by seven protopeptides and color-coded for visual distinction in the schematic diagram. (**E**) The DL model’s effectiveness is validated by demonstrating its ability to accurately extract the core sequences of the 25 oligopeptide drugs from their protopeptides. Subscripts mark the amino acid positions; black subscripts highlight the ground truth of core sequences. Colored sequences (aligned with the colors in panel D) represent the predicted biopeptides near the position of drug core sequences, while the gray sequences depict other biopeptide candidates predicted by the DL model. BP: biological process; MF: molecular function; IDRs: intrinsically disordered regions; PUFs: proteins of unknown function.

Technically, therapeutic oligopeptide discovery fundamentally revolves around two key issues: where to search for therapeutic oligopeptides and how to identify them. Deepeptide employs a two-step pipeline to address the two key issues without the requirement of indication-specific positive data.

For the first issue, where to search, Deepeptide constructs a specialized library of protopeptides with indication-ameliorating-related molecular functions (**Figure 1B**). It is guided by a prior knowledge that oligopeptides are integral components of proteins, from the level of sequence to function^20,21^. Therefore, instead of typically searching therapeutic oligopeptides from peptide library, searching from proteins or their functional regions (that is, protopeptides) with indication-ameliorating-related molecular functions is also a feasible approach. For example, if we aim to discover angiogenic oligopeptides, searching them from protopeptides known to enhance angiogenesis is likely to yield desired results. This step does not need indication-specific positive peptides; instead, it leverages readily available protopeptides obtained through existed computational biology tools^22,23^.

For the second issue, how to identify, Deepeptide solves it by extracting oligopeptides and inferring their function from protopeptides. For oligopeptide extraction, a DL model is developed to identify potential cleavable oligopeptide fragments within protopeptides (**Figure 1C**), guided by a prior knowledge that oligopeptides are typically derived from enzymatic cleavage of protopeptides. Further, we design an algorithm to infer the functions of protopeptide-derived oligopeptides and retain only those with indication-ameliorating-related molecular functions, considering the multi-functionality protopeptides^24^. For example, protopeptides with angiogenic functions may also possess other indication-unrelated functions, performed by different oligopeptides within the protopeptides, while we aim to retain only the angiogenic oligopeptides. The function inference algorithm is based on the principle of functional enrichment, a widely used biological concept^25^, and does not require indication-specific positive data as well.

In this design, there is no need for indication-specific training data, making a generalizable pipeline for various metabolic diseases entirely possible. Furthermore, since the pipeline does not rely on pre-defined target receptors, it can go beyond well-studied targets and even potentially discover leads with novel action mechanisms.

## RESULTS

### Constructing large specialized library composed of protopeptides with indication-ameliorating-related molecular functions

As a two-step pipeline, the first step of Deepeptide tackles the issue of where to search, constructing a large specialized library consisting of protopeptides with indication-ameliorating-related molecular functions, in which oligopeptide candidates are included as sub-fragments. The library construction involves two parts: determining disease indication-ameliorating-related biological processes and molecular functions, and acquiring protopeptides with these molecular functions.

Taking obesity, a disease indication caused by chronic energy imbalance, as an example, its hallmark features include abnormalities in lipid metabolism, such as increased fat synthesis and reduced fat breakdown^26,27^. Therefore, the first step in constructing the specialized library is identifying biological processes and molecular functions ameliorating lipid metabolism. Specifically, we identify 17 biological processes that positively regulate lipid metabolism and retain 1,607 proteins associated with these processes (**Table S1**). These proteins are enriched in 86 molecular functions, of which we select 13 with positive regulatory effects on lipid metabolism (**Table S1**). For example, we included biological processes such as the ‘positive regulation of lipid metabolic process (GO:0045834)’ and ‘positive regulation of lipid catabolic process (GO:0050996)’, as well as molecular functions like ‘triglyceride lipase activity (GO:0004806)’ and ‘lipoprotein lipase activity (GO:0004465)’.

The second part of library construction involves identifying protopeptides with the lipid metabolism-related molecular functions to construct the large specialized library, following the dictum ‘bigger is better’^28^. Intuitively, secreted proteins with functions of interest might appear as natural candidates for protopeptides. However, their limited number and multifunctional nature make them less suitable for constructing a large and specialized library^29,30^. Instead, protein functional sequences, which represent specific regions within proteins, offer significant advantages in terms of both quantity and functional specificity. We therefore investigated which type of functional sequences is most suitable to be protopeptides, based on three key metrics forming the foundation of our pipeline. First, ubiquity, defined as the proportion of sequence lengths across the protein universe, ensures scalability for large library construction. Second, functional diversity, quantified by the number of indication-ameliorating-related functions associated with each sequence type, guarantees broad coverage across diseases. Third, annotation depth, assessed at the molecular function level, connects biological processes, molecular functions, functional regions, and functional oligopeptides.

Among various protein functional sequences, such as localization signal sequences, structure-related regions, and conserved regions, we ultimately select intrinsically disordered regions (IDRs) as the protopeptides for constructing the large specialized library due to their superior performance across all three metrics (**Table S2**). Furthermore, IDRs play a critical role in mediating protein-protein interactions, making them highly relevant for drug discovery^31,32^. For example, in our previous study, we identified an osteogenic pentapeptide within IDRs that mediates the α5-integrin signaling pathway^33^. These characteristics highlight the suitability of IDRs as the foundation for constructing a large, specialized library.

With the determination of lipid metabolism-related molecular functions and the selection of IDRs as protopeptides, we proceeded to identify IDRs with these molecular functions to construct the library. To achieve this, we developed molecular function fingerprint (MFF)-based prediction models using FAIDR^34^, which demonstrates favourable performance in identifying IDRs with specialized functions. Leveraging this model, we identified a large number of functional IDRs from the protein universe. Notably, the protein universe is primarily composed of Proteins of Unknown Function (PUFs)^35,36^, and therefore functional IDRs in the constructed large specialized library are primarily derived from protein ‘dark matter’, particularly PUFs. This protein ‘dark matter’-broadened large specialized library enables Deepeptide to discover novel candidates in terms of both sequence and action mechanism.

### Developing general screening algorithm using deep learning

The second step of Deepeptide solves the issue of how to identify by developing a general screening algorithm, involving two parts: extracting oligopeptide candidates and inferring their functions from functional protopeptides. As aforementioned, the modeling principle is guided by prior knowledge that endogenous biopeptides are primarily cleaved from protopeptides at specific enzyme recognition sites and may retain or perform certain specific functions of the protopeptides.

For the first part, we construct a DL model to extract oligopeptide candidates cleaved from protopeptides. Actually, series of DL models have been developed to identify cleavage sites or extract biopeptides within protopeptides. For example, PeptideLocator^22^ and SignalP^23^ are developed to identify endogenous or signal peptides from protopeptides. These tools utilize the information of cleavage sites and protopeptides as training data and employ the classic named entity recognition framework of natural language processing, where a DL model discerns named entities by learning the patterns that distinguish entities from their context.

Similarly, our DL model also adapts the classic named entity recognition framework^37^ and is developed with three main components (**Figures 1C and S1**): 1) Input Representation: We fine-tuned the state-of-the-art pre-training protein large language model, ESM-2^38^, to integrate global evolutionary information and general semantic patterns of proteins; 2) Context Encoder: A bidirectional Long Short-Term Memory (Bi-LSTM) network^39^ captures context dependencies, crucial for extracting the targeted subsequence (in this case, the oligopeptide) from the larger sequence (the protopeptide); 3) Tag Decoder: We employed a Conditional Random Field (CRF) model to assign tags to each amino acid^39^, indicating its membership as part of the biopeptide.

To train our DL model, we constructed a protopeptide dataset containing endogenous biopeptides and their flanking sequences separated by cleavage sites (**Table S3**). Herein, flanking sequences represent the upstream and downstream amino acid residues adjacent to these biopeptides within their protopeptides, and are distinguished from biopeptides using the cleavage sites as boundaries. These biopeptides are explicitly defined as endogenous in their source databases, collected from UniProt^40^ and 18 publicly available biopeptide databases (**Table S4**). Protopeptides in this dataset are labeled at the amino acid level as biopeptide or flanking sequence, using ‘B’, ‘I’, ‘E’, or ‘O’, indicating the amino acid is in the starting point, internal region, endpoint and outside of a biopeptide. Herein, ‘B’ and ‘E’ indicate cleavage sites and ‘O’ represents flanking sequences. Consequently, the deep learning model is trained to identify biopeptides near hydrolysis cleavage sites in protopeptides, regardless of established action mechanisms or ligand characteristics like amino acid composition.

As the second part, we infer functions of oligopeptide candidates employing their functional enrichment in the library, inspired by the principle of protein function enrichment. We propose that oligopeptides enriched in the library of functional protopeptides with specific molecular functions are likely to possess corresponding molecular functions. Accordingly, we devise indicators of ‘enrichment significance’ and ‘function score’ to evaluate the potential of oligopeptides to possess the corresponding molecular functions. As a result, we constructed a general DL-based algorithm, capable of screening oligopeptide candidates with indication-ameliorating-related molecular functions, without the need for indication-specific training data.

### Validating the DL model on marketed oligopeptide drugs

To evaluate the ability of our DL model to extract biopeptides from protopeptides, we validated its performance using 25 marketed oligopeptide drugs across diverse indications, derived from the seven core sequences of their parent proteins (**Figure 1D and Table S5**). For rigor, the DL model was retrained on a revised dataset, excluding the core sequences of these 25 marketed drugs and their parent proteins from the original training set. Results showed that our DL model accurately extracted five core sequences without any errors and pinpointed two core sequences with a shift of only 1 or 3 amino acids (**Figure 1E**). This suggests that the DL model has effectively learned the general splitting pattern between biopeptides and flanking sequences. Moreover, the model’s precision in extracting only a limited number of oligopeptides—approximately three biopeptide candidates per 1,000 amino acids—significantly reduces the size of candidate pools (**Figure 1E**). This reduction offers a distinct advantage by decreasing both screening time and discovery costs in biopeptide research. Overall, these results highlight the efficiency and reliability of our DL model for extracting potential biopeptides from protopeptides.

### Illustrative process of Deepeptide for therapeutic oligopeptide discovery

We applied Deepeptide on five representative disease indications of metabolism, endocrinology, and tissue regeneration: lipid metabolism, osteogenesis, glucose metabolism, angiogenesis, and anti-angiogenesis.

As an illustrative example, we here describe the two-step pipeline for angiogenic oligopeptide discovery (**Figure 2A, Table S6**). For the first step, we constructed a specialized library composed of protopeptides with indication-ameliorating-related molecular functions (**Figure 2B**). The process involved: 1) Using ‘angiogenesis’ as a keyword of disease indication, we collected 28 biological processes and retained 20 biological processes that positively regulate angiogenesis. For example, positive regulation of sprouting angiogenesis (GO:1903672) (**Table S1**). Then, for each of these biological processes, we retained only proteins that promote angiogenesis, resulting in 453 proteins with angiogenic function. Furthermore, we analyzed these 453 proteins and found they were enriched in 49 molecular functions, from which we selected 9 molecular functions with positive regulatory effects on angiogenesis. Such as, platelet-derived growth factor receptor binding (GO: 0005161) (**Table S1**). 2) Using FAIDR, we constructed an MFF-based prediction model for these molecular functions (**Figure S2A**) to screen the functional IDRs from the protein universe, obtaining a specialized library composed of 35,936 angiogenesis-related functional IDRs. This process could be easily reusable to each of these indications initiated with their respective indication keywords — ‘lipid metabolism’ for lipid metabolism, ‘bone formation’ for osteogenesis, ‘insulin, GLP-1, and α-amylase’ for glucose metabolism, and ‘angiogenesis’ for both angiogenesis and anti-angiogenesis.

**Figure 2.**
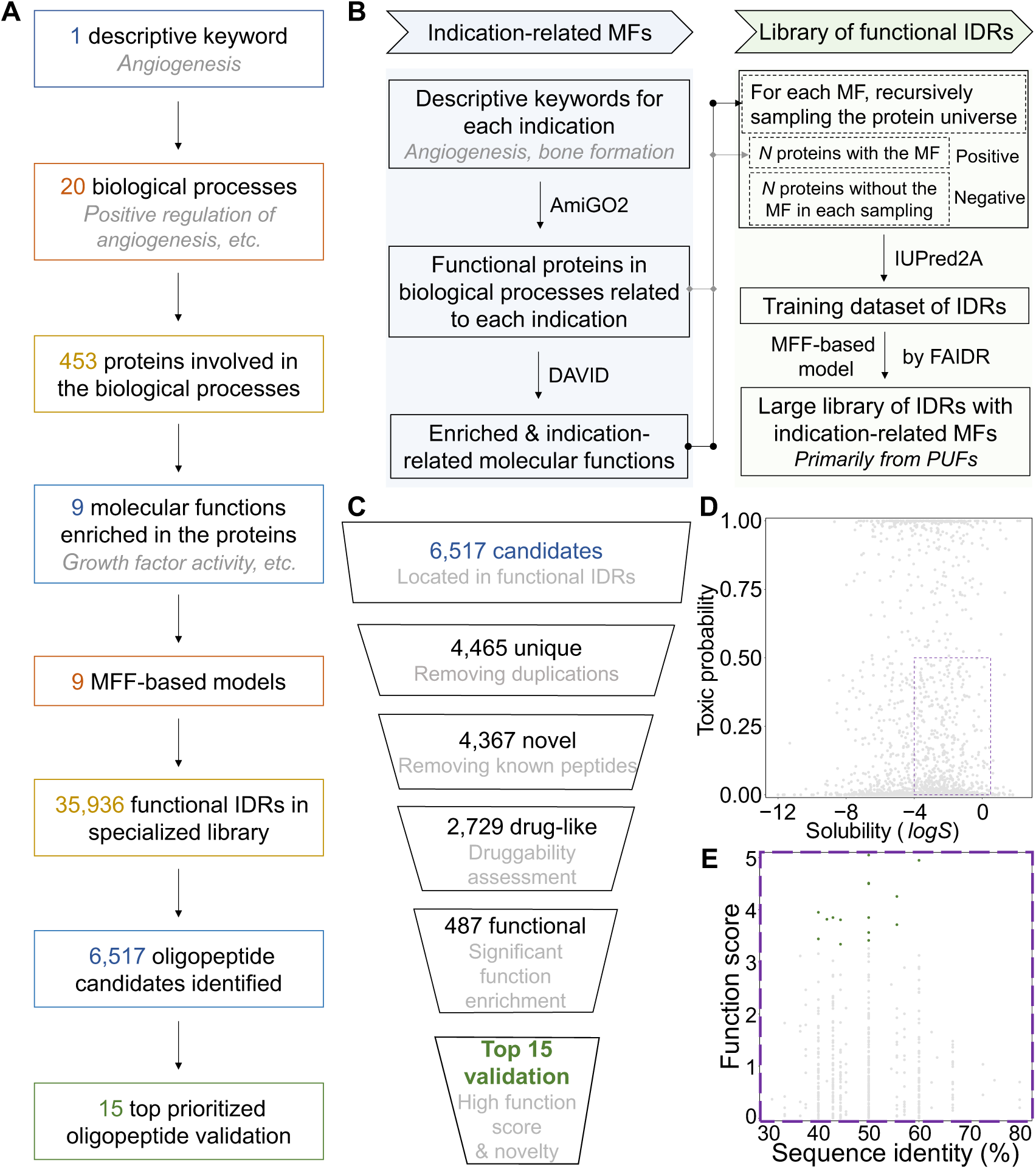
Utilizing Deepeptide to discover therapeutic oligopeptides, illustrated with the indication of angiogenesis. **(A**) A workflow outlines the process of discovering therapeutic oligopeptide candidates, using angiogenesis as a specific indication example. (**B**) The process of constructing a specialized library for the interested disease indication is shown. The process consists of two steps: determining the indication-ameliorating-related biological processes and molecular functions, and screening IDRs with these indication-ameliorating-related molecular functions using the molecular function fingerprint (MFF)-based prediction model. (**C)** A prioritization process narrows down 6,517 oligopeptide candidates identified by the DL model to the top-15 candidates for wet-lab validation. **(D)** A scatter plot shows the distribution of solubility (x-axis) against toxicity (y-axis) for oligopeptide candidates. Candidates within the purple dashed box are deemed favorable in terms of solubility (*log S* value between -4 and 0.5) and safety (toxic probability less than 0.5) and are selected for further prioritization (as shown in panel E). **(E)** A scatter plot depicts the highest sequence identity to oligopeptides in the training dataset (x-axis) against function score (y-axis) of the oligopeptide candidates. The top 15 ranked oligopeptides, selected based on function score, are highlighted for subsequent wet-lab validation.

For the second step, we extracted the oligopeptide candidates by our DL model and prioritized them. The process involved (**Figure 2C**): 1) Extraction of 6,517 oligopeptide candidates from the specialized library with our DL model (**Figure S3A and S3B**). 2) Elimination of duplicated oligopeptides identified in different IDRs, which left 4,465 biopeptide candidates. 3) Elimination of oligopeptides overlapped with known peptides of the training dataset, which filtered out 2.2% of them and left 4,367 candidates (**Figure S3C**). 4) Further refinement through solubility and toxicity screenings to retain oligopeptides with favourable druggability, which left 2,729 candidates (**Figure 2D**). 5) Retention of candidates that were significantly enriched in the specialized library, with selection narrowed to 487 candidates. 6) Selection of the top-15 candidates for wet laboratory validation based on their function score. These candidates exhibit relatively low sequence identity (49.5% in average) with the known peptides in the training dataset (**Figure 2E**). The low overlap ratio and sequence identity indicate the capacity of Deepeptide to identify novel sequences.

### AP7 with comparable potency to VEGF for angiogenesis

For the indication of angiogenesis, we assessed fifteen therapeutic oligopeptide candidates through a series of *in vitro* validations, including wound healing (scratch assay), cell viability (CCK-8 assay), and tube formation assays. Of these candidates, seven demonstrated significantly enhanced angiogenic potential compared to the untreated control, enhancing angiogenesis by promoting cell migration rather than cell proliferation (**Figure S4**), which minimizes the risk of tumor formation while ensuring angiogenesis^41,42^. Five candidates exhibited angiogenic potency comparable to that of VEGF, and the most efficacious candidate, AP7, was selected for further *in vivo* validations.

We assessed the angiogenic effects of AP7 using an excisional wound splinting mouse model^43^ (**Figure 3A**), considering the crucial role of blood vessel formation in wound healing^44^. Wound closure rates, measured on days 3, 7, and 12 post-injury (**Figure 3B, 3C**), revealed that wounds in the AP7-treated group closed significantly faster than those in the PBS group, showing rates comparable to those in the VEGF-treated group. Additionally, no adverse reactions were observed in any of the mice throughout the 12-day experimental period.

**Figure 3.**
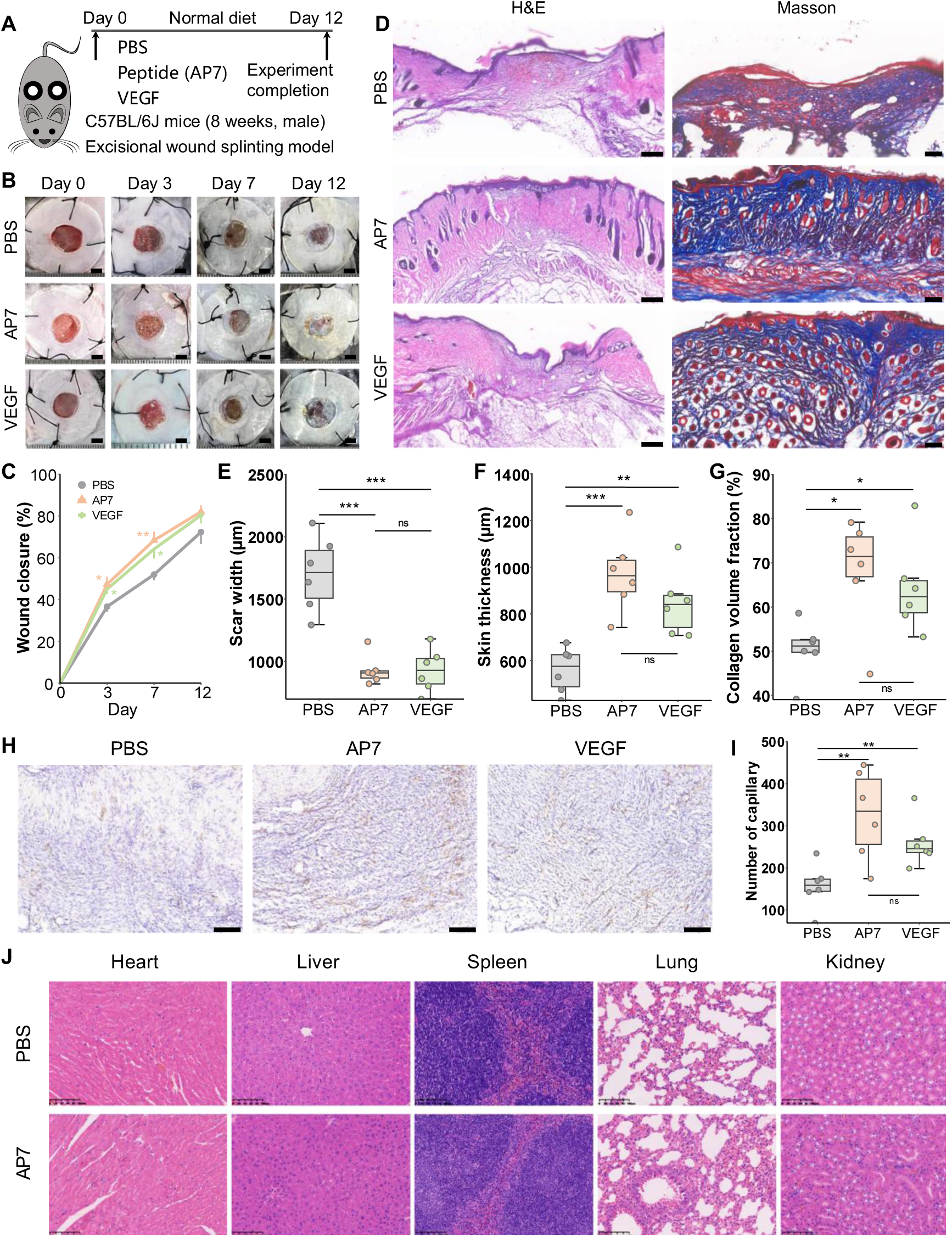
AP7 with comparable potency to VEGF for angiogenesis. **(A)** A schematic represents the *in vivo* experiment assessing the angiogenic effects of AP7 using an excisional wound splinting model in mice. Mice are divided into three groups, receiving daily topical treatments of either PBS, AP7 (2 μg/mL), or VEGF-A 145 (0.25 μg/mL) respectively, during the 12-day experiment. **(B)** Representative images show wounds in mice treated with PBS, AP7, or VEGF-A 145 at the indicated time points. Scale bar: 2 mm. **(C)** A graph illustrates the rate of wound closure at indicated time points; N = 7 per group. Data are expressed as means ± SEM. **(D)** Histological analysis of full-thickness wounds using H&E and Masson’s trichrome staining 12 days post-injury. Scale bars: 200 μm for H&E, 100 μm for Masson’s trichrome. **(E**-**G)** Quantitative assessments of scar width, skin thickness, and collagen volume fraction in full-thickness wounds 12 days post-injury; N = 6 per group. **(H)** Immunohistochemistry staining for CD31 in wounds 12 days post-injury. Scale bar: 100 μm. **(I)** Quantification of capillaries from CD31 immunohistochemistry staining; N = 6 per group. **J**, Representative H&E-stained images of heart, liver, spleen, lung, and kidney tissues collected from mice at the completion of the experiment. Scale bar: 100 μm. Significance markers: \**P* < 0.05, \*\**P* < 0.01, \*\*\**P* < 0.001; ns: not significant.

H&E staining of tissue samples from day 12 post-injury revealed that the AP7-treated group exhibited enhanced re-epithelialization and dermis regeneration with reduced scar formation, compared to the PBS group, and showed comparable results to the VEGF group (**Figure 3D-F**). Masson’s trichrome staining indicated more extensive collagen deposition and better-organized collagen fibers in the AP7-treated group, again comparable to the VEGF group (**Figure 3D, 3G**). Immunohistochemistry staining for the vascular endothelial marker CD31 showed significantly increased neovascularization in the AP7-treated group relative to the PBS group, at levels similar to those in the VEGF group (**Figure 3H, 3I**). Additionally, no significant histopathological alterations were observed in the heart, spleen, lung, or kidney tissues of mice in AP7-treated group compared to the PBS group (**Figure 3J**). Collectively, these findings demonstrate the capacity of Deepeptide to discover potent and safe oligopeptides significantly stimulating angiogenesis and accelerating wound healing, with efficacy on par with VEGF and novel action mechanism.

### TP6 as a bifunctional oligopeptide ameliorates hyperlipidemia and obesity

Obesity is driven by chronic energy imbalance, characterized by abnormalities in lipid metabolism that contribute to both adipose tissue accumulation and dysregulated lipid levels in circulation^27^. One hallmark of obesity is the expansion of white adipose tissue, primarily driven by the differentiation of precursor cells into adipocytes^26^. Additionally, hyperlipidemia—characterized by elevated blood lipid levels—is a key metabolic feature of obesity and a major contributor to its associated complications^45^. Thus, targeting lipid metabolism, including inhibiting adipogenesis and reducing circulating lipid levels, represents a promising strategy for managing obesity and its comorbidities^46^. In this context, fifteen oligopeptides for lipid metabolism identified by Deepeptide were initially evaluated for their anti-adipogenic and lipid-lowering activities through *in vitro* validation assays, including Oil Red O staining and expression analysis on thermogenesis-related genes (*Ucp1*, *Pgc1a*) and lipid synthesis-related gene (*Pparg*). Of these, nine demonstrated comparable efficacy, while one exhibited superior efficacy compared to GLP-1, Exendin-4, Simvastin, and Celastrol (**Figure S5**). The most efficacious candidate, TP6, demonstrated promising anti-adipogenic and lipid-lowering activity in the initial screening of therapeutic candidates and was selected for subsequent *in vivo* validations.

We evaluated the treatment effects of TP6 on HFD-induced hyperlipidemia and obesity in HFD mice. Remarkably, intraperitoneal injection of TP6 at 4 mg/kg resulted in a significant decrease in body weight gain compared to that in HFD controls over 11 weeks of HFD exposure (**Figure 4A-C**), while no significant difference in food intake was observed among the groups (**Figure 4D**). Moreover, we repeated this assay with two positive control drugs, Exendin-4 for obesity and Simvastin for hyperlipidemia. As results (**Figure S6A**), there were almost no significant changes in the weight of mice in different treatment groups, although the Exenatide-4-treated mice tended to lose more weight during the first three weeks, the weight gradually caught up over the later period. This suggests that TP6, as a naked peptide, is comparable to the GLP-1 receptor agonists Exenatide-4 for weight loss. And, compared with HFD group, high levels of lipids in the blood (TG, TC, and LDL-c) and liver (TG and TC) were significantly reduced by TP6 treatment, with comparable efficacy to the Simvastatin group (**Figure 4E, 4F, Figure S6B**). Consistent with these findings, H&E and Oil Red O staining revealed that the pathological lesions, such as massive accumulation of lipid droplets in the liver and increased fat cell size caused by HFD feeding, could be obviously reversed by TP6 treatment (**Figure 4G**). Additionally, no significant histopathological abnormalities were observed in the heart, spleen, lung, or kidney tissues of mice treated with TP6 compared to the NC and HFD group (**Figure S6C**), suggesting that TP6 exhibits favorable safety at the therapeutic dose. Taken together, we conclude that TP6 has both lipid-lowering and weight-reducing effects and is comparable to the first-line drugs.

**Figure 4.**
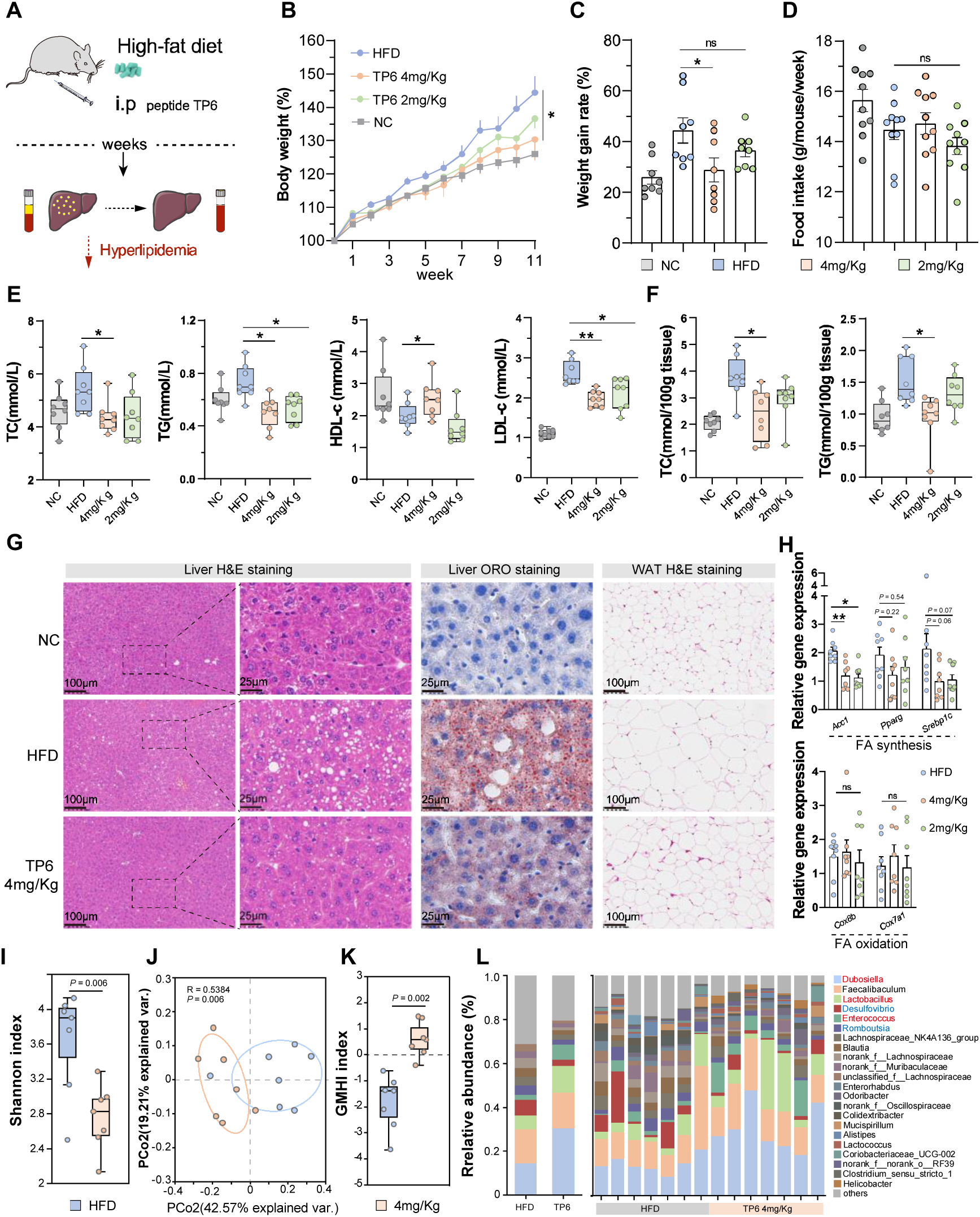
TP6 as a bifunctional oligopeptide ameliorates hyperlipidemia and obesity. **(A)** A schematic illustrates the animal experiment assessing the treatment effect of TP6 on HFD-induce hyperlipidemia and obesity in mice; i.p: intraperitoneal injection. Mice are divided into four groups and fed a standard normal chow diet or 60% fat diet for 10 weeks; the TP groups receive intraperitoneal injection of different doses of TP6, and saline serves as a parallel control (NC); N = 8 per group. **(B**-**C)** Percentage of body weight gained measured over time in treated mice; N = 8 per group. **(D)** Weekly measurements of food intake; N = 8 per group. **(E)** Serum lipid profiles, including triglycerides (TG), total cholesterol (TC), high-density lipoprotein cholesterol (HDL-c), and low-density lipoprotein cholesterol (LDL-c); N = 8 per group. **(F)** Lipid contents of TG and TC in liver tissues. **(G)** Representative histological images of liver and fat tissues stained with H&E and Oil red O staining under different treatments. Scale bars = 100 or 25 μm as applicable. **(H)** Gene expression analysis in liver tissue; N = 8 per group. **(I)** Shannon index of fecal microbiota in HFD and TP6-treated mice. **(J)** PCA of fecal microbiota in HFD and TP6-treated mice. The significance of two separated clusters is measured with the Adonis test. **(K)** Gut Microbiota Health Index (GMHI) of fecal microbiota. **(L)** Fecal microbiota composition in HFD and TP6-treated mice at genus levels. Significantly increased and decreased genus levels are marked in red and blue letters, respectively. For panels **B**, **C**, **D,** and **H**, data are expressed as means ± SEM. Significance markers: \**P* < 0.05, \*\**P* < 0.01, \*\*\**P* < 0.001. ns, not significant. ORO: Oil red O staining; NC: negative control; WAT: white adipose tissues.

To determine whether TP6 affects fat metabolism, we investigated the expression of enzymes involved in lipid synthesis and oxidation using a real-time PCR system. Our results showed that TP6 decreased the relative mRNA levels of *acetyl-CoA carboxylase 1* (*Acc1*)^47^, *peroxisome proliferator-activated receptor gamma* (*Pparg*)^48^, and *sterol regulatory element-binding protein 1C* (*Srebp1c*)^49^ (**Figure 4H**), which lead to cellular lipid accumulation. Notably, TP6 treatment did not alter the activity of enzymes involved in fat oxidation, such as *Cytochrome C Oxidase Subunit 7A1* (*Cox7a1*) and *Cytochrome C Oxidase Subunit 8B* (*Cox8b*) (**Figure 4H**). These findings indicate that TP6, as a representative oligopeptide screened for lipid metabolism, may inhibit fat accumulation in a dose-dependent manner by regulating the expression of lipid synthesis factors, such as *Acc1* and *Srebp1c*, but not lipid oxidation, thereby ameliorating hyperlipidemia and liver steatosis.

The gut microbiota, a key contributor to metabolic function, has been linked to the development of hyperlipidemia and obesity^50,51^. To further understand the role of gut microbiota in alleviating hyperlipidemia and obesity through TP6 treatment, we investigated the composition of the intestine-dwelling microbiome using 16S rRNA gene amplicon sequencing. While α-diversity showed a significant decrease, inter-group β-diversity, as visualized by Principal Coordinates Analysis (PCoA), revealed four distinct clusters depending on the treatment, suggesting differences in gut microbiota composition in response to HFD and TP6 treatment (**Figure 4I, J**). Furthermore, we found that the TP6 group had a much higher gut microbiome health index (GMHI), a robust index for assessing health status based on species-level taxonomic characteristics of gut microbiome samples^52^ (**Figure 4K**). In addition, compared with the HFD mice, TP6 treatment significantly decreased the levels of *Desulfovibrio* and *Romboutsia*, which were positively correlated with obesity^53^, and increased levels of *Dubosiella*, *Lactobacillus*, and *Enterococcus* (**Figure 4L**). These results demonstrate that TP6 treatment can improve the intestinal bacterial disorders caused by an HFD.

Together, these results underscore the ability of Deepeptide to identify potent and safe therapeutic oligopeptides that achieve efficacy in ameliorating lipid metabolism comparable to the first-line drugs, even exhibiting novel action mechanism.

### Discovery of potent oligopeptides for osteogenesis, glucose metabolism, and anti-angiogenesis

Furthermore, we applied Deepeptide to three additional indications: osteogenesis, glucose metabolism, and anti-angiogenesis (**Figure 5A**). *In vitro* validations were conducted to assess the oligopeptide candidates using MC3T3-E1, HUVEC, and HepG2 cell lines to evaluate osteogenic, glucose metabolic, and anti-angiogenic activities, respectively.

**Figure 5.**
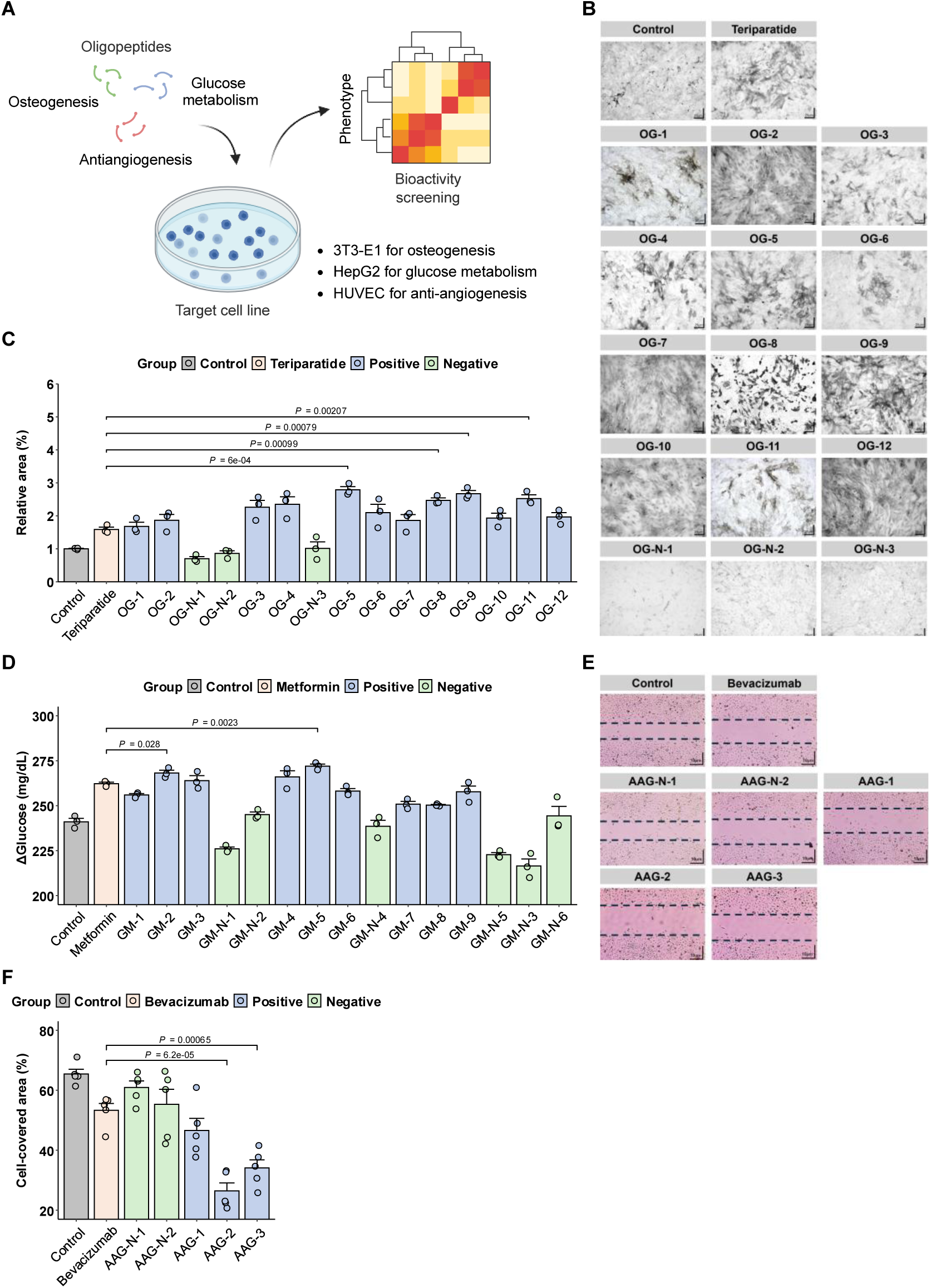
Validation of oligopeptides for indications of osteogenesis, anti-angiogenesis, and glucose metabolism. **(A)** An overview of *in vitro* assays conducted at 0.1 ug/mL concentration for osteogenesis (B and C), glucose metabolism (D) and anti-angiogenesis (E and F), with each experiment performed in triplicate. **(B)** Representative images of ALP-stained MC3T3-E1 cells after osteogenic induction with oligopeptide treatment (Scale bar = 25 μm). **(C)** Quantification of osteoblast cell area after oligopeptide treatment is performed using ImageJ, calculated as the percentage of ALP-stained osteoblast coverage relative to the total area. **(D)** Glucose consumption in HepG2 cells after oligopeptide treatment is quantified by measuring the change in absorbance at 630 nm, corresponding to the blue-green Schiff base formed through the reaction of glucose with the o-toluidine reagent. **(E)** Scratch assay is performed on HUVEC cells treated with oligopeptides (Scale bar = 10 μm). **(F)** Quantification of cell coverage after oligopeptide treatment is calculated as the ratio of the area covered by cell migration (difference between the initial scratch area and the post-migration area) to the initial scratch area. Data are presented as means ± SEM. Significance markers: *\*P* < 0.05, *\*\*P* < 0.01, *\*\*\*P* < 0.001, *\*\*\*P* < 0.0001.

For the osteogenesis indication, among the fifteen evaluated oligopeptide candidates, 80% (12/15) demonstrated significantly enhanced osteogenic differentiation compared to controls, with eight showing comparable efficacy and four exceeding that of Teriparatide (**Figure 5B, C**). In the glucose metabolism indication, among the fifteen candidates assessed, 60% (9/15) significantly improved intracellular glucose metabolism, with six showing comparable efficacy and two surpassing the potency of Metformin (**Figure 5D**). For the indication of anti-angiogenesis, among the five candidates validated, 60% (3/5) significantly reduced cell migration, with three matching and two surpassing the potency of Bevacizumab group (**Figure 5E, F**). Additionally, 68% (27/40) of the positive oligopeptides across all five indications were derived from PUFs, demonstrating the substantial potential of leveraging protein ‘dark matter’ for oligopeptide lead discovery.

## DISCUSSION

In this study, we presented a generalizable pipeline for therapeutic oligopeptide discovery applicable to various metabolic diseases, based on the understanding of the relationships among disease indications, biological processes and molecular functions. The identified novel oligopeptide candidates have demonstrated promising potential as oligopeptide leads, exhibiting potency comparable to the first-line drugs by novel action mechanisms.

The well-known street-light effect^54^ also occurs in the field of drug discovery—the vast majority of research concentrates on a very small number of indications, while most indications receive little attention^5,55^. AI-driven drug discovery holds the promise of alleviating this inequality, with its key aspiration of developing a broadly generalizable pipeline^28^. Here, Deepeptide made its endeavor for metabolic diseases. Encouragingly, with the increasingly accumulated protein sequences and the emerging of increasingly accurate bioinformatics tools for annotating the functions of protein sequences^29,56^, it will be progressively easier for Deepeptide to build specialized libraries and identify therapeutic oligopeptides, and thus be applied for more unmet indications.

Going beyond the extensively studied targets or action mechanisms with AI-driven drug discovery is highly anticipated by the scientific and industrial community^17^, presenting higher requirements for both library construction and screening algorithms. From the perspective of library construction, it is essential to integrate a large number of diverse and previously unexplored candidates. Recent studies have demonstrated the potential of identifying novel antimicrobial peptides from immunity-unrelated proteins in the human proteome^10^, ancient proteomes^57^, and microbiomes^9,58^. These findings highlight the vast and largely untapped protein ‘dark matter’ as a rich resource for discovering novel lead candidates beyond the scope of widely studied mechanisms of action. Similarly, this study introduced PUFs to construct a large specialized library comprising diverse and unexplored oligopeptide candidates. In terms of screening algorithms, effective models should operate independently of pre-defined ligand characteristics or well-known receptor structures. In peptide drug discovery, AI-based screening algorithms often employed classification or regression models tailored to specific indications^1,5^. While these models can be effective, they are highly dependent on large datasets and are inherently constrained by the pre-defined characteristics of ligands^5,7,59^. To overcome the limitation, Deepeptide establishes a novel framework for the identification and function inference of biopeptides. Jointly, these breakthroughs in both library construction and screening algorithm make Deepeptide more feasible to identify potent oligopeptides novel in sequences and action mechanisms.

In summary, our study proposed a DL-based pipeline, capable of discovering potent and novel oligopeptide leads for various metabolism-related diseases. This highlights the feasibility of constructing a reliable AI-based application by leveraging fundamental biological knowledge, even when positive data are extremely scarce. We anticipate that these results will significantly accelerate the discovery of therapeutic oligopeptide leads for numerous clinical indications in the near future.

### Limitations of study

The limitations of this study can be summarized into three main aspects. First, Deepeptide is not suitable for discovering therapeutic peptides for indications lacking well-characterized biological processes, as its ‘disease indications — biological processes — molecular functions’ pipeline depends on the availability of known indication-ameliorating-related biological processes. Second, Deepeptide performs more effectively in identifying short biopeptides, such as oligopeptides, since its enrichment-based approach may be less effective for long biopeptides due to their lower occurrence frequency. Lastly, Deepeptide is unable to account for modifications such as acetylation or pegylation due to the inherent limitations of the deep learning model employed.

## STAR METHODS

### KEY RESOURCES TABLE

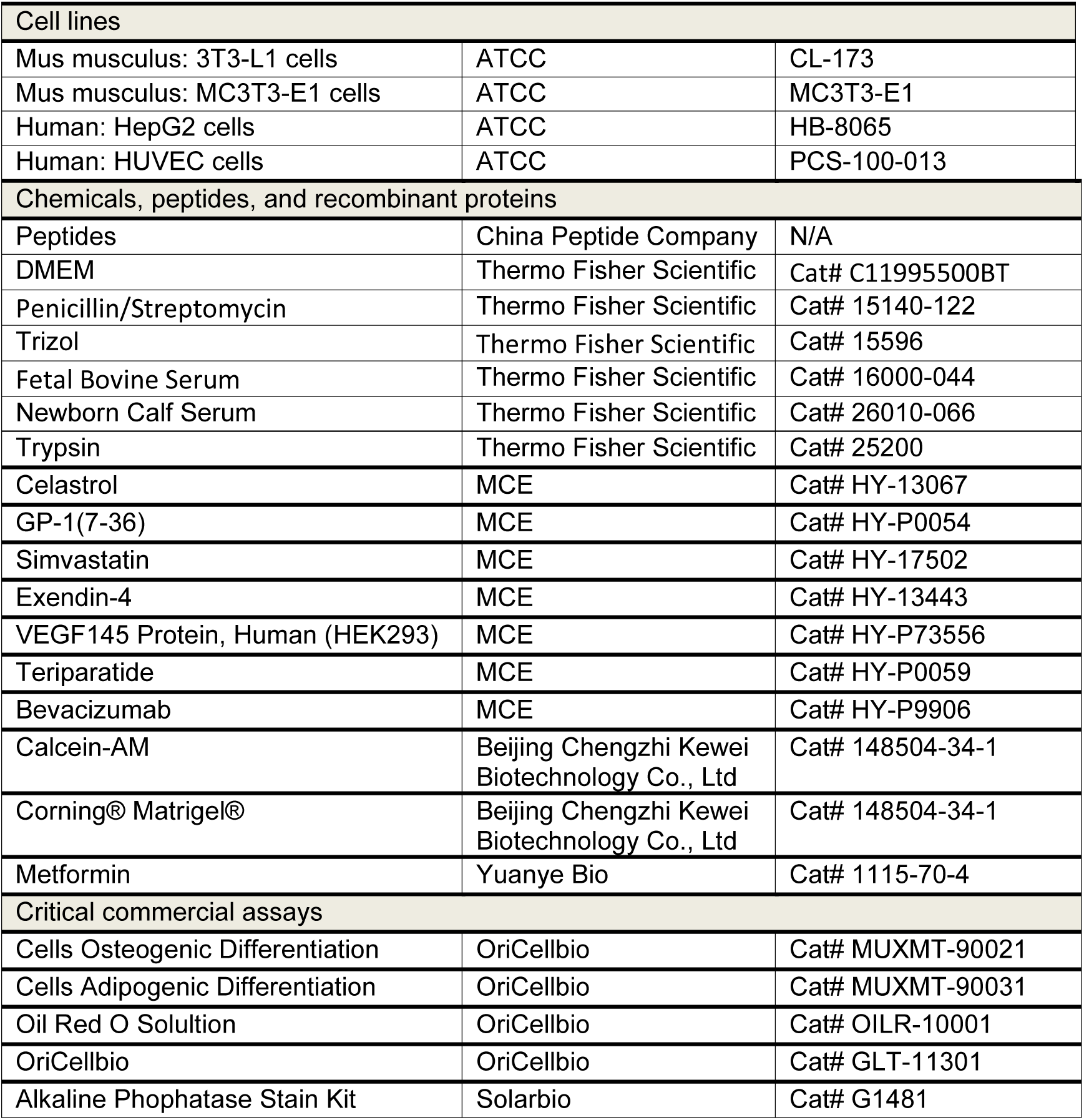
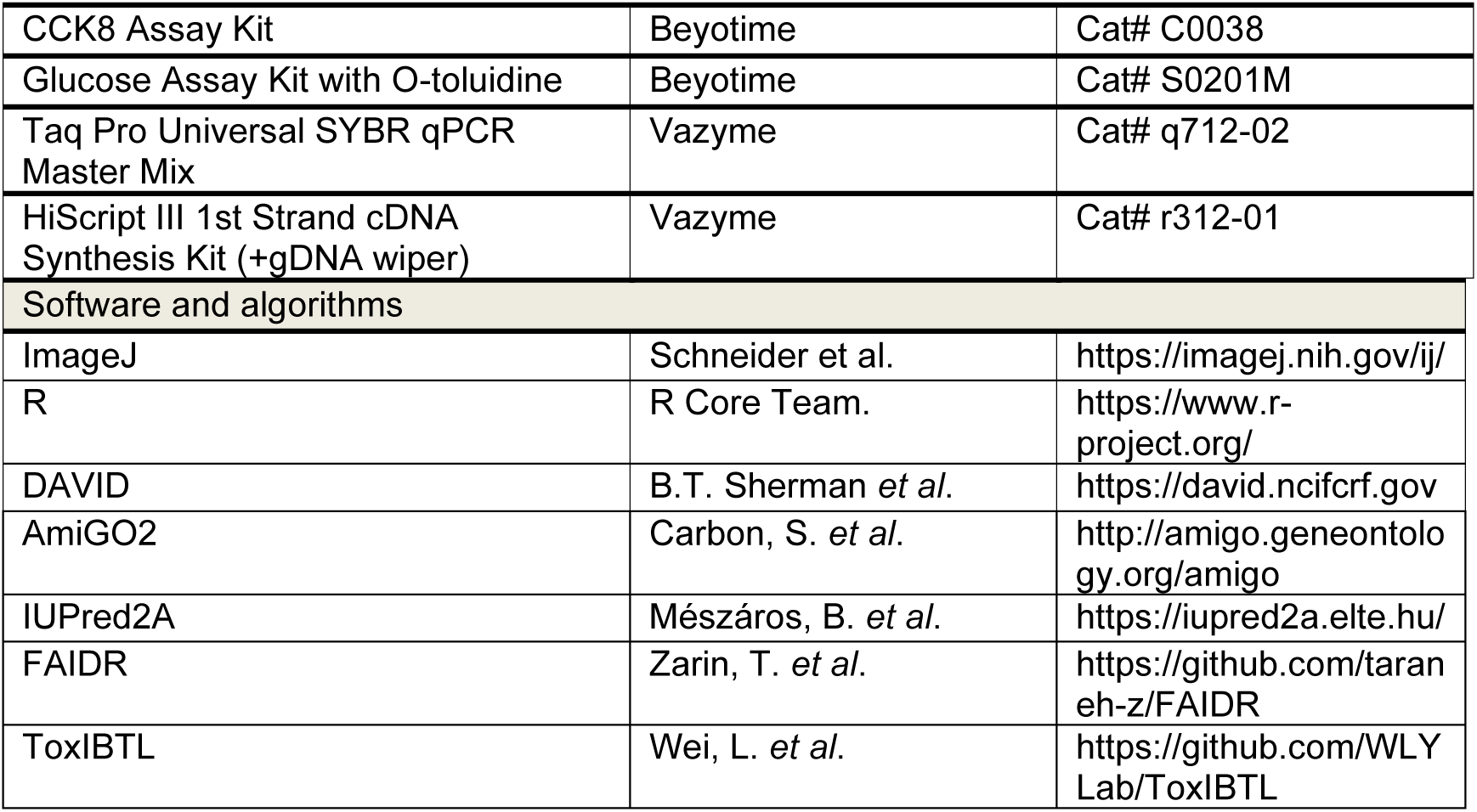

## RESOURCE AVAILABILITY

### Lead contact

Further information and requests for resources and reagents should be directed to and will be fulfilled by the lead contact, Yong-Biao Zhang.

### Materials availability

All the materials generated in this study are available upon reasonable request to the lead contact. This study did not generate new unique reagents.

### Data and code availability

The data underlying each numeric display item is available in the ‘**Data S1** – Source Data’ excel file. The custom code for the general screening algorithm can be found at https://github.com/xiaobaichuan/Deepeptide. Any additional information required to reanalyze the data reported in this paper is available from the lead contact upon request.

## EXPERIMENTAL MODEL AND SUBJECT DETAILS

### Animals and experiments

All C57BL/6J mice, sourced from SPF Biotechnology (Beijing, China), were accommodated in a pathogen-free facility. Groups of 4-6 animals were housed per cage with standard bedding and environmental enrichment. The facility maintained controlled temperature and humidity conditions and operated on a 12-hour light/dark cycle. Mice had unrestricted access to standard laboratory chow and water. Except as noted for specialized experiments, all animals were maintained on this standard diet. In experiments with an excisional wound splinting model, 8-week-old male C57BL/6J mice were selected.

In two independent repeated experiments with HFD mice, 7-week-old male C57BL/6J mice were switched to a high-fat diet (60 kcal% fat, D12492; Research Diets, Inc., New Brunswick, NJ) for 7 weeks or 11 weeks. During this period, their body and food weights were recorded weekly. The oligopeptide TP6 was administered daily via intraperitoneal injection. In HFD mice experiments compared with first-line drug, TP6, Exendin-4, Simvastatin are administered at 4 mg/kg, 10 µg/kg and 1.5mg/kg, respectively.

### Excisional wound splinting model

The excisional wound splinting model involves immobilizing full-thickness skin wounds using circular silicone splints to minimize measurement errors from skin contraction. These splints have a 6 mm inner diameter and a 15 mm outer diameter. Following intraperitoneal anesthesia, the mice’s dorsal hair is removed, and two 5 mm diameter full-thickness skin wounds are created using a skin punch. The wounds are encircled with the silicone rings using adhesive and further secured with sutures. Gauze and bandages are then applied to protect the wounds and splints from external contamination. During the 12-day experiment, mice with these wounds received daily topical treatments of either PBS, AP7 (2 μg/mL), or VEGF-A 145 (0.25 μg/mL). On day 12, the mice were euthanized. Digital images of the wounds were taken on days 0, 3, 7, and 12, and analyzed using ImageJ to calculate wound healing rates. The healing rate was determined using the formula: healing rate (%) = (1 - [residual wound area at each time point] / [initial wound area]) × 100%.

### Cell lines

3T3-L1 cells were cultured in High Glucose Dulbecco’s Modified Eagle Medium (DMEM-H) supplemented with 10% calf serum (CS), under conditions of 5% CO2 and 37°C (**key resources table**). For differentiation, cells were initially treated with DMEM-H containing 10% CS, 0.5 mM 3-isobutyl-1-methylxanthine, 1 mM dexamethasone, and 1 μg/mL insulin for two days, followed by two days in DMEM-H with 10% CS and 1 μg/mL insulin. Subsequently, they were maintained in DMEM-H with 10% CS until significant lipid droplet formation was observed. HUVEC cells were cultured in DMEM-H with 10% Fetal Bovine Serum (FBS); HepG2 cells in DMEM supplemented with 10% FBS and GlutaMAX; and MC3T3-E1 cells in α-MEM with 10% FBS, followed by treatment with peptides at specified concentrations in osteogenic induction medium. All cell lines were sourced from the Cell Resource Center, Peking Union Medical College, which is the headquarters of NSTI-BMCR (National Science & Technology infrastructure National BioMedical Cell-Line Resource) (**key resources table**).

## METHOD DETAILS

### Definitions of protopeptide, biopeptide, oligopeptide and therapeutic peptide

A protopeptide refers to a precursor, which can be a complete protein or a protein fragment, from which biopeptides are derived. In this study, we specifically utilized functional IDRs as protopeptides due to their relevance in protein-protein interactions and therapeutic potential.

A biopeptide is defined as a peptide with specific biological activity. In this study, biopeptides were identified from protopeptides using our deep learning model. These biopeptides are bioactive fragments with therapeutic or regulatory potential.

An oligopeptide, as defined here, is a biopeptide consisting of 4 to 10 amino acids.

This lower bound (4 amino acids) corresponds to the shortest known therapeutic oligopeptides derived from proteins in marketed drugs^4^. While the upper limit for oligopeptide length varies in the literature (ranging from 10 to 20 amino acids)^60–62^, we chose 10 as the cutoff to maintain specificity and align with the widely accepted principle that “shorter is better” for certain therapeutic applications^8,63^.

A therapeutic peptide is defined as a biopeptide with the capacity to heal or treat a disease or disorder^4^. Thus, in this study, a therapeutic peptide refers to a biopeptide with demonstrated therapeutic properties.

### Training dataset construction

To train our deep learning model, we compiled a dataset comprising 7,028 protein-derived endogenous biopeptides (4 to 50 amino acids) mapped to 7,894 protopeptides (precursor proteins) (**Table S3**). These data were obtained from UniProt (https://www.uniprot.org/; Release 2024_06) and 18 publicly available biopeptide databases (**Table S4**). Specifically, biopeptides and paired information of cleavage sites and protopeptides were obtained from the PTM/Processing section of UniProt or directly extracted from the 18 biopeptide databases, excluding viral or toxic entries.

For the 7,894 protopeptides, those exceeding 1,024 amino acids in length were sequentially split into 1,024-amino-acid segments to comply with the input length limitation of the DL model. This segmentation process resulted in a total of 9,175 protopeptides. Each amino acid within the protopeptides was subsequently labeled according to its positional information within the protopeptides using the BIEO scheme. In this scheme, amino acids are assigned one of four tags: ‘B’, ‘I’, ‘E’, or ‘O’, indicating the amino acid is in the starting point, internal region, endpoint and outside of a biopeptide, respectively.

When employed to implement model evaluation, the dataset was evenly split into five subsets using GraphPart^64^, with redundancy between any two subsets limited to below 30%. For each fold, the model was retrained on four subsets, and the remaining subset was used to assess its performance in predicting cleavage sites. As results, the average recall metric of five-fold cross validation was respectively 0.64, 0.69, 0.73, and 0.76 within tolerance thresholds of 0, 1, 2, and 3 around the true cleavage sites, and the average precision metric was respectively 0.75, 0.82, 0.84, and 0.86 within tolerance thresholds of 0, 1, 2, and 3 around the true cleavage sites. These findings suggest that the deep learning model has learned to identify biopeptides in regions proximate to cleavage sites.

### Architecture of the deep learning model

#### Input

We take protopeptides with a length of no more than 1,024 amino acids as input for the deep learning model. The length limitation of 1,024 amino acids is set by the large language model ESM, as 99% of all sequences in MGnify90 and 96.7% of sequences in UniParc contain fewer than 1,024 amino acids^38,65^.

#### Input representation

For constructing the module of input representation, we fine-tuned the pre-trained Transformer model, ESM-2, a protein large language model developed by Meta AI research^38,65^. ESM-2 learns the semantic patterns of billions of protein sequences through self-supervised learning and demonstrates strong performance and transferability across tasks, such as predicting solvent accessibility, secondary structure, and structural disorder^66^. For each input, ESM-2 returns its feature tensor with dimension of sequence_length*1,280, that is, each amino acid in the sequence is encoded as a feature vector of length 1,280.

#### Context encoder

To encode context information of protopeptides, we employed Bi-LSTM^39^. Bi-LSTM effectively learns the implicit splitting patterns between biopeptides and their flanking sequences. Here, num_layers = 2 and hidden_size = 256.

#### Tag decoder

We used CRF^39^ as the decoder to translate the output of the Bi-LSTM into tags for amino acids, indicating whether they belong to biopeptides or not. As the most commonly used decoder, CRF captures the intrinsic correlations among tags, leading to improved performance compared to decoding tags independently.

#### Output

We utilized four types of tags to represent the labels of amino acids. The tags ‘B’, ‘I’, and ‘E’ indicate the beginning, inner, and end of biopeptides, respectively, while ‘O’ signifies that amino acid is outside of any biopeptide.

### Training process of the deep learning model

#### Fine-tuning

We fine-tuned ESM-2 by the procedure described in the paper of ESM^65^ and its sample code (https://github.com/facebookresearch/esm). Specifically, we fine-tuned the pre-trained Transformer model by optimizing the parameters of F(x) with the supervised objective, where F(x) is the model combination of ‘ESM-2, Bi-LSTM, CRF’, and supervised objective is the loss of sequence tagging.

#### Loss function

The loss function utilizes the negative log-likelihood of the predicted and true labels, calculated by the CRF function in torchcrf module.

#### Optimizing

We optimized the model using AdamW with a learning rate of 3e-5.

### Collecting core sequences of marketed oligopeptide drugs

The core sequence of a marketed drug, as defined in this study, refers to the peptide consisting exclusively of the 20 natural amino acids that retains the drug’s therapeutic properties. In practice, marketed peptide drugs are often chemically modified to enhance pharmacokinetic properties, such as stability or bioavailability. These modifications frequently involve residues beyond the standard amino acids, making the publicly disclosed sequences unsuitable for direct use in computational models. For example, the publicly available sequence of Icatibant is represented as ‘rRPXGXSXR,’ where ‘X’ indicates chemically modified amino acids introduced during drug optimization. Since our deep learning models are specifically designed to process only natural amino acids, the original, unmodified peptide (referred to as the ‘naked peptide’) was used as the core sequence.

The marketed peptide drugs included in this study were compiled from PepTherDia^4^, comprising a total of 112 marketed peptide drugs. Among these, 72 are classified as oligopeptide drugs, with 25 having publicly available sequence information that can be directly obtained from PepTherDia or acquired through its recorded references in DrugBank^67^ and PubChem^68^.

To extract the core sequences of these oligopeptide drugs, we employed a two-step methodology combining sequence mapping and literature review (**Table S5**). For 19 drugs, core sequences were successfully identified by mapping their sequences onto parent protein sequences. For the remaining drugs, such as Bremelanotide and Eptifibatide, core sequences were determined through extensive manual literature review.

The sequence mapping process consisted of two steps. First, information about the parent protein and the ‘masked’ sequence of the oligopeptide drug was obtained from PepTherDia. For example, Icatibant is derived from bradykinin-related peptides, with a masked sequence of ‘rRPXGXSXR’. The second step involved mapping the ‘masked’ sequence to its corresponding region within the parent protein. In the case of Icatibant, the masked sequence ‘rRPXGXSXR’ was mapped to ‘RRPPGFSPFR’ within the bradykinin-related peptide sequence, identifying it as the core sequence. This process yielded the core sequences of 25 marketed oligopeptide drugs, derived from seven distinct parent proteins (**Figure 1D**).

### Selecting the indications for therapeutic oligopeptide lead discovery

Currently, peptide drugs primarily target indications related to endocrinology, metabolism, oncology, and the central nervous system^3^. In this study, considering their broad relevance and potential clinical utility (**Figure S2B**), we selected five representative indications for therapeutic oligopeptide lead discovery: angiogenesis, lipid metabolism, osteogenesis, glucose metabolism, and anti-angiogenesis.

### Constructing the specialized library composed of functional IDRs

To construct the specialized library composed of functional IDRs for the selected indications – angiogenesis, lipid metabolism, osteogenesis, glucose metabolism, and anti-angiogenesis – we initiated the process with descriptive keywords of each indication. These keywords were ‘angiogenesis’ (used for both angiogenesis and anti-angiogenesis), ‘lipid metabolism’ (for lipid metabolism), ‘bone formation’ (for osteogenesis), and ‘insulin, GLP-1, and α-amylase’ (for glucose metabolism). Taking ‘angiogenesis’ as a case study, we detail the following processes, which were similarly applied to the other indications using their respective keywords.

#### Determining the indication-ameliorating-related molecular functions

To determine molecular functions related to the ‘angiogenesis’ indication, we first submitted the keyword of ‘angiogenesis’ to AmiGO2 (http://amigo.geneontology.org/amigo)^69^ to retrieve the relevant biological processes (**Table S1**). Next, for each biological process, we retained only those functional proteins involved in promoting angiogenesis filtered in the operational panel ‘GO class’ of AmiGO2, and further excluded proteins from species other than *Homo sapiens*, *Mus musculus*, and *Rattus norvegicus* in the operational panel ‘Organism’ of AmiGO2. Taking ‘regulation of cell migration involved in sprouting angiogenesis’ (GO:0090049) as an example, proteins involved in ‘positive regulation of biological process’ are retained, while those involved in ‘negative regulation of biological process’ are discarded. Finally, we conducted a gene enrichment analysis for the retained proteins (termed as functional proteins) using DAVID v2022q1 (https://david.ncifcrf.gov/)^70^, focusing on the molecular functions precisely promoting angiogenesis and linking to at least 10 proteins (considering the accuracy of subsequent MFF-based model construction) (**Table S1**).

#### MFF-based model construction

To construct the MFF-based prediction model, we first utilized IUPred2A (https://iupred2a.elte.hu/)^71^ to predict IDRs in all proteins across the protein universe. For each indication-ameliorating-related molecular function, we then compiled a training dataset consisting of IDRs derived from functional proteins exhibiting the specific molecular function as the positive dataset. An equal number of proteins unrelated to these functional proteins were randomly selected from the protein universe, of which IDRs form the negative training dataset. This approach assumes that proteins other than the functional proteins randomly selected from the protein universe (usually accessed from UniProt) are unlikely to share the specific biological functions of interest. Finally, we utilized FAIDR (https://github.com/taraneh-z/FAIDR) to build the MFF-based prediction model (**Figure S2A**).

#### Functional IDRs identification

Using the MFF-based prediction model, we identified functional IDRs with the indication-ameliorating-related molecular functions within the positive and negative datasets. To maintain balanced training datasets and ensure robust model performance, we employed a no-replacement sampling strategy to process all IDRs in the protein universe across multiple training epochs. For each epoch, the model was retrained with a newly constructed dataset, allowing comprehensive evaluation of the IDRs in relation to the specific molecular functions.

As a proof-of-concept, the protein universe in this study refers to all 384,651 proteins of the above three species (*Homo sapiens*, *Mus musculus*, and *Rattus norvegicus*) collected from UniProt. Clearly, researchers can construct a larger specialized library by screening the functional IDRs from a bigger set of protein sequences, such as the ancient proteomes, microbiomes, etc.

### Prioritization of oligopeptide candidates

To prioritize oligopeptide candidates identified from functional IDRs using the DL model, we considered the following joint attributes:

#### Enrichment significance

For each oligopeptide *i*, its functional enrichment significance is evaluated in two steps. First, we use the MFF-based model, to create datasets of equal size for both functional and non-functional IDRs. We then use two arrays to separately record the occurrences of oligopeptide *i* in each IDRs containing oligopeptide *i* for the two datasets, and maintain the same length for both arrays (i.e., the shorter array is zero-padded to match the length of the longer array), and perform a one-sided *t*-test for the two arrays. Oligopeptides with higher mean value of occurrences in the functional IDRs alongside with an adjusted *P*-value(*i*) < 0.05 advance to the next step; others are set aside. Second, we consider the entire sequence length (*N*) and the total occurrences (*Mi*) of oligopeptide *i* in the protein universe as background parameters. Comparatively, we treat the entire sequence length (*n*) and the occurrences (*mi*) of oligopeptide *i* in the functional IDRs dataset as sampling parameters. Clearly, this forms a hypergeometric distribution, from which we can determine the significance of functional enrichment for oligopeptide *i* as follows:

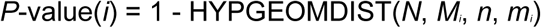

Here, HYPGEOMDIST refers to the value of cumulative distribution function based on the hypergeometric distribution. If herein adjusted *P*-value(i) < 0.05, oligopeptide *i* is considered significantly enriched in the functional IDRs dataset. Here, all original *P*-values are adjusted using the Benjamini and Hochberg (BH) method for multiple testing correction.

#### Function score

The function score is grounded in the principle that higher occurrences of oligopeptides in functional protopeptides indicate greater functional significance. However, to counteract the inherent bias where shorter oligopeptides typically have higher occurrences, we have formulated the function score as follows:

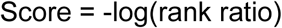

Where, the ‘rank ratio’ for each oligopeptide is determined by its frequency rank percentage within its length-specific sequence cluster. For instance, if a pentapeptide ranks 20th in frequency among 100 pentapeptides, its rank ratio is 0.2. This approach allows us to normalize for length bias. The use of the logarithmic function in the score calculation amplifies the distinctions in function scores, accommodating the generally uniform distribution of rank ratio values across different oligopeptides. Notably, this function score ranks peptide candidates by their relative potential to possess specific molecular functions, enabling experimental prioritization rather than binary classification of YES/NO. In practice, we recommend researchers to select the top *N* candidates based on the function score for wet lab verification, depending on the throughput of their laboratory platform. In this study, we selected the candidates of top 15 for the indications of angiogenesis, lipid metabolism, osteogenesis and glucose metabolism, and top 5 for the indication of anti-angiogenesis.

#### Sequence novelty

To assess sequence novelty, we evaluate an oligopeptide’s homology against those in our training set, using the highest sequence identity as a measure. Lower identity scores indicate greater novelty. This is done using the Needleman–Wunsch algorithm, implemented via the ‘needleall’ function in Jemboss software package (v1.5)^72^ with default parameter settings.

#### Solubility

*S*olubility is a fundamental pharmacochemical property, relevant to a drug’s ADME (absorption, distribution, metabolism, excretion) attributes. We evaluate solubility of oligopeptides by the *logS* (aqueous solubility value) indicator using mordred (v1.2.1a1)^73^, and retain those with values ranging from -4 to 0.5.

#### Toxicity

Early elimination of toxic oligopeptides is essential. We utilize ToxIBTL^74^ to predict toxicity, retaining only oligopeptides identified as non-toxic.

In summary, the prioritization process serves two main purposes, both aimed at enhancing the robustness of candidate selection. First, the function evaluation step filters out potential false positives by assessing the functional relevance of oligopeptide candidates to the target indication. Candidates with low enrichment significance or poor function scores, indicating a reduced likelihood of possessing the desired biological activity, are deprioritized for further validation. Second, the process refines candidate selection by evaluating druggability attributes, including solubility and toxicity. These metrics help identify candidates with higher potential for succeeding progression in the subsequent pipeline. Importantly, druggability evaluation operates independently of function evaluation, allowing users to tailor the prioritization process to their specific experimental needs and preferences.

### Oligopeptide synthesis

We synthesized the oligopeptides through China Peptides Co., Ltd., ensuring a purity level of over 98% for those used in animal experiments (200 mg per peptide) and above 90% for *in vitro* validations (10 mg per peptide). For experimental use, we prepared sterile aqueous stock solutions of these peptides, subsequently storing them at -80 °C and utilizing them within a two-week timeframe. All the oligopeptides tested in this study were unmodified naked peptides, with their sequences provided in **Table S7**.

### Controls for *in vitro* validations

For *in vitro* validations, we utilized distinct positive control drugs for each indication, as detailed in the **key resources table**. VEGF-A 145 at 30 ng/mL served as the control for angiogenesis, Celastrol at 0.3 μmol/L for lipid metabolism, Teriparatide at 10 nmol/L for osteogenesis, Metformin at 0.5 mmol/L for glucose metabolism, and Bevacizumab at 0.5 μg/mL for anti-angiogenesis. All scrambled controls for oligopeptides in the five indications were deposited in **Figure S7**. In addition, GLP-1, Exendin-4, and Simvastatin were also included as controls in the experiment for the validation of TP-6 (LM-7) for lipid metabolism (**Figure S5C**).

### Oil Red O staining

*In vitro* adipogenic-induced 3T3-L1 cells were first washed with PBS, then fixed for 10 minutes using 4% (v/v) paraformaldehyde. After a 20-second wash with staining wash solution, the cells were incubated with Oil Red O solution (OILR-10001, Oricellbio) for 10-20 minutes and subsequently washed twice more with the staining wash solution. Images of the stained cells were captured using a MF31-M microscope (Mshot, China). Quantitative analysis of these images was performed using ImageJ software v1.8.0.345 (National Institutes of Health, Bethesda, Maryland, USA).

### Cell scratch assay

For the cell migratory assays, HUVEC cells were grown in DMEM-H supplemented with 10% FBS to full confluence. A scratch was then created in the cell monolayer using a sterile 200-µL pipette tip. Post-scratching, the cells were washed twice with PBS to clear debris and detached cells, and subsequently, serum-free medium was introduced to the wells. The plate was incubated at 37°C and 5% CO2, with images captured at 0, 24, and 48 hours using an MF31-M microscope (Mshot, China). The extent of cell coverage across the scratch was quantified using ImageJ software v1.8.0.345 (National Institutes of Health, Bethesda, Maryland, USA).

### Tube Formation Assay

In the tube formation assay, HUVEC cells were prepared at 80% confluence, then diluted to a density of 1×10^5^ cells/mL and seeded into a 96-well plate. Cells were treated with peptides at a concentration of 0.2 µg/mL, followed by incubation as per standard cell culture conditions. After an 8-hour culture period, the cells were washed with DPBS and stained with Calcein AM for 40 minutes at 37°C. Tube formation was observed using a fluorescence microscope, set to an excitation wavelength of 494 nm and an emission wavelength of 515 nm. Quantitative analysis of the tube structures, including measurements of tube length, coverage area, and the number of loops and nodes, was conducted using the Angiogenesis Analyzer plugin in ImageJ.

### ALP and Glucose staining

For ALP staining, we used a kit from Beyotime (P0321S). Osteogenic-induced MC3T3-E1 cells were first washed with PBS and then fixed with 4% (v/v) paraformaldehyde for 10 minutes. These cells were subsequently immersed in an ALP staining buffer for 10 minutes at room temperature. After removing the staining buffer, cells were washed twice with PBS. Images of the stained cells were captured using an MF31-M microscope (MF31-M, Mshot, China), with quantitative analysis performed using ImageJ software v1.8.0.345 (National Institutes of Health, Bethesda, Maryland, USA).

For Glucose consumption, we used the Beyotime kit (S0201M). HepG2 cells were washed twice with PBS, lysed using cell lysis buffer, and centrifuged at 12,000 × g for 5 minutes. The supernatant was mixed with the Glucose Assay Reagent (o-toluidine reagent), which reacts with glucose to form a blue-green Schiff base with a maximum absorption wavelength of 630 nm. The mixture was incubated at 95°C for 8 minutes, cooled to 4°C, and the absorbance at 630 nm was measured using an iMark™ Microplate Absorbance Reader (1681130 BIO-RAD, USA). Glucose consumption in the samples was quantified by analyzing the differential absorbance at 630 nm measured before and after peptide stimulation.

### Quantitative real-time PCR assay

For the *in vitro* validations across the five indications, total RNA was extracted from harvested cells using Trizol reagent (Thermo, Cat# 15596). cDNA synthesis was performed using the HiScript DNA Synthesis Kit with gDNA wiper (Vazyme, Cat# r312-01), and quantitative PCR (qPCR) was conducted with the SYBR qPCR Master Mix kit (Vazyme, Cat# q712-02). Relative mRNA levels were normalized to the housekeeping gene GAPDH in accordance with MIQE guidelines. Primer sequences, designed using Oligo6 software v6.62 (Molecular Biology Insights, Colorado, USA), are detailed in **Source data**. All the qPCR results of five indications were deposited in **Figure S8**.

For animal experiments in HFD mice, total RNA was extracted with Trizol reagent (Invitrogen) according to the manufacturer’s instructions as previously described^75^. We used 1 μg of total RNA for reverse transcription with an enzyme from Yeasen (Shanghai, China). RT-qPCR was carried out using SYBR Green PCR master mix (Yeasen) on an ABI7500 real-time PCR system (Applied Biosystems). The primers, listed in **Source data**, were used to calculate gene expression as a percentage relative to GAPDH.

### Morphology and histology

For angiogenesis assessment, skin tissues from the wound sites of the PBS, VEGF, and AP7-treated groups were collected on day 12. These samples were fixed overnight in 4% paraformaldehyde (PFA) solution at 4 °C and subsequently sectioned to a thickness of 5 µm for frozen sections. We performed Hematoxylin and Eosin (H&E) and Masson’s trichrome staining on these sections as per the manufacturer’s instructions. For immunohistochemical analysis, sections were first incubated with rabbit anti-CD31 primary antibody (#77699, 1:200, CST) overnight at 4 °C, followed by a 30-minute room temperature incubation with horseradish peroxidase (HRP)-conjugated secondary antibodies (1:400, ZSGB-BIO). DAB (3,3’-Diaminobenzidine) staining was then applied to visualize CD31-positive endothelial cells, indicating vascular structures within the tissue. Whole-tissue slides were scanned using a Pannoramic MIDI microscope slide scanner (3DHISTECH). Images at 10× and 20× magnification were analyzed using CaseViewer_2.3 software. Scar width, skin thickness, collagen volume fraction, and vessel densities were quantified using ImageJ.

For the indication of lipid metabolism, liver and epididymal white adipose tissues (WAT) were promptly harvested. The liver WAT was then divided, with one portion immediately stored at -80°C and the other fixed in 4% paraformaldehyde (PFA) solution for 24 hours, followed by paraffin embedding. Tissue sections of 5 µm were prepared and subjected to Hematoxylin and Eosin (H&E) and Oil Red O staining, in accordance with the manufacturer’s instructions. Whole-tissue slides were scanned at 40× magnification using a Pannoramic MIDI microscope slide scanner (3DHISTECH). Images at 40× and 80× magnifications were analyzed using CaseViewer_2.3 software.

### Biochemical test

Biochemical analyses for triglycerides (TG), total cholesterol (TC), high-density lipoprotein cholesterol (HDL-C), and low-density lipoprotein cholesterol (LDL-C) in plasma and liver samples were conducted using assay kits from Nanjing Jiancheng Bioengineering Institute (Nanjing, Jiangsu, China). These tests were performed in strict accordance with the manufacturer’s provided protocol.

### DNA isolation, and 16S rRNA sequencing

Bacterial DNA was extracted from fecal samples using the QIAamp-DNA Mini Stool Kit (QIAGEN, CA, USA), following the manufacturer’s instructions. The extracted DNA was quantified by agarose gel electrophoresis and spectrophotometrically using a NanoDrop One system (Thermo Scientific, Waltham, USA), then stored at -80 °C for future analysis. For 16S rRNA amplicon sequencing, the V3 and V4 regions of the bacterial 16S rRNA gene were amplified using modified region-specific primers (341F-805R), 2-5 μL of extracted DNA, and sequencing primers with Illumina paired-end adapters, utilizing Q5 Hot Start Polymerase (NEB, Ipswich, MA). The intestinal flora’s abundance and diversity in mice were assessed through Illumina HiSeq sequencing (I-Sanger, Beijing, China). Bioinformatics analysis of the sequences was performed using the QIIME software package, as previously described^75^. Sequences with 97% identity were grouped into the same operational taxonomic units (OTUs) using an open reference picking strategy based on the Greengenes database. PCoA was carried out with QIIME, based on unweighted UniFrac distances, and inter-group comparisons were made using Wilcoxon rank-sum tests.

### Statistics

Statistical analyses were conducted using R software, version 4.2.2. Except as noted for specialized analysis, significance testing was carried out using the one-way ANOVA. Correlation analyses were performed based on the Spearman’s rho statistic. Statistical significance was denoted as follows: \**P* < 0.05, \*\**P* < 0.01, \*\*\**P* < 0.001, \*\*\*\**P* < 0.0001, and ‘ns’ indicating non-significant.

## Supporting information

Supplementary Figure 1-8 (termed as Figure S1-S8)

Supplementary Table 1-7 (termed as Table S1-S7)

## ACKNOWLEDGMENTS

This research was supported by grants from the National Natural Science Foundation of China (32470644, 82171844 to Y.-B.Z.) and China Postdoctoral Science Foundation (GZB20230196 to Z.L.).

## AUTHOR CONTRIBUTIONS

Y.-B.Z. and B.X. conceptualized and designed the overall project. The architecture of Deepeptide was established by B.X., C.M., and Z.F., who also undertook the data analysis. The *in vitro* and *in vivo* experiments were conducted by Z.L., X.W., H.Z., B.X., R.L., H.S., R.H., D.L., X.T. and Z.S. The manuscript was initially drafted and revised by B.X., H.Z., Z.L., and Y.-B.Z. Z.M., Y.Z., S.W., L.W., J.H., J.L., C.L., H.H., L.W. and J.W. provided crucial data analysis and insightful project suggestions.

## DECLARATION OF INTERESTS

The authors declare that they have no competing interests.

## Notes

### Competing Interest Statement

The authors have declared no competing interest.

## REFERENCES

1. Basith, S., Manavalan, B., Hwan Shin, T., and Lee, G. (2020). Machine intelligence in peptide therapeutics: a next-generation tool for rapid disease screening. Med. Res. Rev. 40, 1276–1314.

2. Henninot, A., Collins, J.C., and Nuss, J.M. (2018). The current state of peptide drug discovery: back to the future? J. Med. Chem. 61, 1382–1414.

3. Muttenthaler, M., King, G.F., Adams, D.J., and Alewood, P.F. (2021). Trends in peptide drug discovery. Nat. Rev. Drug Discov. 20, 309–325.

4. D Aloisio, V., Dognini, P., Hutcheon, G.A., and Coxon, C.R. (2021). PepTherDia: Database and structural composition analysis of approved peptide therapeutics and diagnostics. Drug Discov. Today 26, 1409–1419.

5. Chen, Z., Wang, R., Guo, J., and Wang, X. (2024). The role and future prospects of artificial intelligence algorithms in peptide drug development. Biomed. Pharmacother. 175, 116709.

6. Jiménez-Luna, J., Grisoni, F., and Schneider, G. (2020). Drug discovery with explainable artificial intelligence. Nat. Mach. Intell. 2, 573–584.

7. Paul, D., Sanap, G., Shenoy, S., Kalyane, D., Kalia, K., and Tekade, R.K. (2021). Artificial intelligence in drug discovery and development. Drug Discov. Today 26, 80.

8. Huang, J., Xu, Y., Xue, Y., Huang, Y., Li, X., Chen, X., Xu, Y., Zhang, D., Zhang, P., and Zhao, J. (2023). Identification of potent antimicrobial peptides via a machine-learning pipeline that mines the entire space of peptide sequences. Nat. Biomed. Eng. 7, 797–810.

9. Ma, Y., Guo, Z., Xia, B., Zhang, Y., Liu, X., Yu, Y., Tang, N., Tong, X., Wang, M., and Ye, X. (2022). Identification of antimicrobial peptides from the human gut microbiome using deep learning. Nat. Biotechnol. 40, 921–931.

10. Torres, M.D., Melo, M.C., Flowers, L., Crescenzi, O., Notomista, E., and de la Fuente-Nunez, C. (2022). Mining for encrypted peptide antibiotics in the human proteome. Nat. Biomed. Eng. 6, 67–75.

11. Agrawal, P., Bhagat, D., Mahalwal, M., Sharma, N., and Raghava, G.P. (2021). AntiCP 2.0: an updated model for predicting anticancer peptides. Brief. Bioinform. 22, bbaa153.

12. Lv, Z., Cui, F., Zou, Q., Zhang, L., and Xu, L. (2021). Anticancer peptides prediction with deep representation learning features. Brief. Bioinform. 22, bbab008.

13. Rao, B., Zhou, C., Zhang, G., Su, R., and Wei, L. (2020). ACPred-Fuse: fusing multi-view information improves the prediction of anticancer peptides. Brief. Bioinform. 21, 1846–1855.

14. Fu, X., Cai, L., Zeng, X., and Zou, Q. (2020). StackCPPred: a stacking and pairwise energy content-based prediction of cell-penetrating peptides and their uptake efficiency. Bioinformatics 36, 3028–3034.

15. Kardani, K., and Bolhassani, A. (2021). Cppsite 2.0: an available database of experimentally validated cell-penetrating peptides predicting their secondary and tertiary structures. J. Mol. Biol. 433, 166703.

16. Manavalan, B., and Patra, M.C. (2022). MLCPP 2.0: an updated cell-penetrating peptides and their uptake efficiency predictor. J. Mol. Biol. 434, 167604.

17. Sethi, A., and Rathi, B. (2024). Artificial intelligence in drug discovery: a mirage or an oasis? In 103994.

18. Akbarian, M., Khani, A., Eghbalpour, S., and Uversky, V.N. (2022). Bioactive peptides: Synthesis, sources, applications, and proposed mechanisms of action. International journal of molecular sciences 23, 1445.

19. Damodaran, S. (2008). Amino acids, peptides and proteins. Fennema’s food chemistry 4, 425–439.

20. Akbarian, M., Khani, A., Eghbalpour, S., and Uversky, V.N. (2022). Bioactive peptides: Synthesis, sources, applications, and proposed mechanisms of action. International journal of molecular sciences 23, 1445.

21. Damodaran, S. (2008). Amino acids, peptides and proteins. Fennema’s food chemistry 4, 425–439.

22. Bonetta, R., and Valentino, G. (2020). Machine learning techniques for protein function prediction. Proteins: Structure, Function, and Bioinformatics 88, 397–413.

23. Zarin, T., Strome, B., Peng, G., Pritišanac, I., Forman-Kay, J.D., and Moses, A.M. (2021). Identifying molecular features that are associated with biological function of intrinsically disordered protein regions. Elife 10, e60220.

24. Zanzoni, A., Ribeiro, D.M., and Brun, C. (2019). Understanding protein multifunctionality: from short linear motifs to cellular functions. Cell. Mol. Life Sci. 76, 4407–4412.

25. Huang, D.W., Sherman, B.T., and Lempicki, R.A. (2009). Bioinformatics enrichment tools: paths toward the comprehensive functional analysis of large gene lists. Nucleic. Acids. Res. 37, 1–13.

26. Lin, X., and Li, H. (2021). Obesity: epidemiology, pathophysiology, and therapeutics. Front. Endocrinol. 12, 706978.

27. Xia, W., Veeragandham, P., Cao, Y., Xu, Y., Rhyne, T.E., Qian, J., Hung, C., Zhao, P., Jones, Y., and Gao, H. (2024). Obesity causes mitochondrial fragmentation and dysfunction in white adipocytes due to RalA activation. Nat. Metab. 6, 273–289.

28. Sadybekov, A.V., and Katritch, V. (2023). Computational approaches streamlining drug discovery. Nature 616, 673–685.

29. Bonetta, R., and Valentino, G. (2020). Machine learning techniques for protein function prediction. Proteins: Structure, Function, and Bioinformatics 88, 397–413.

30. Zanzoni, A., Ribeiro, D.M., and Brun, C. (2019). Understanding protein multifunctionality: from short linear motifs to cellular functions. Cell. Mol. Life Sci. 76, 4407–4412.

31. Blundell, T.L., Gupta, M.N., and Hasnain, S.E. (2020). Intrinsic disorder in proteins: Relevance to protein assemblies, drug design and host-pathogen interactions. Progress in Biophysics and Molecular Biology 156, 34–42.

32. Uversky, V.N. (2020). Intrinsically disordered proteins: targets for the future? Structural Biology in Drug Discovery: Methods, Techniques, and Practices, 587–612.

33. Cai, M., Xiao, B., Jin, F., Xu, X., Hua, Y., Li, J., Niu, P., Liu, M., Wu, J., and Yue, R. (2022). Generation of functional oligopeptides that promote osteogenesis based on unsupervised deep learning of protein IDRs. Bone Res. 10, 23.

34. Zarin, T., Strome, B., Peng, G., Pritišanac, I., Forman-Kay, J.D., and Moses, A.M. (2021). Identifying molecular features that are associated with biological function of intrinsically disordered protein regions. Elife 10, e60220.

35. Horan, K., Jang, C., Bailey-Serres, J., Mittler, R., Shelton, C., Harper, J.F., Zhu, J., Cushman, J.C., Gollery, M., and Girke, T. (2008). Annotating genes of known and unknown function by large-scale coexpression analysis. Plant Physiol. 147, 41–57.

36. Wang, W., Shuai, Y., Yang, Q., Zhang, F., Zeng, M., and Li, M. (2024). A comprehensive computational benchmark for evaluating deep learning-based protein function prediction approaches. Brief. Bioinform. 25, bbae050.

37. Li, J., Sun, A., Han, J., and Li, C. (2020). A survey on deep learning for named entity recognition. IEEE Trans. Knowl. Data Eng. 34, 50–70.

38. Lin, Z., Akin, H., Rao, R., Hie, B., Zhu, Z., Lu, W., Smetanin, N., Verkuil, R., Kabeli, O., and Shmueli, Y. (2023). Evolutionary-scale prediction of atomic-level protein structure with a language model. Science 379, 1123–1130.

39. Huang, Z., Xu, W., and Yu, K. (2015). Bidirectional LSTM-CRF models for sequence tagging. arXiv preprint arXiv:1508.01991.

40. 2024). UniProt: the Universal protein knowledgebase in 2025. Nucleic. Acids. Res., gkae1010.

41. Lugano, R., Ramachandran, M., and Dimberg, A. (2020). Tumor angiogenesis: causes, consequences, challenges and opportunities. Cell. Mol. Life Sci. 77, 1745–1770.

42. Saman, H., Raza, S.S., Uddin, S., and Rasul, K. (2020). Inducing angiogenesis, a key step in cancer vascularization, and treatment approaches. Cancers 12, 1172.

43. Wang, X., Ge, J., Tredget, E.E., and Wu, Y. (2013). The mouse excisional wound splinting model, including applications for stem cell transplantation. Nat. Protoc. 8, 302–309.

44. Tonnesen, M.G., Feng, X., and Clark, R.A. (2000). Angiogenesis in wound healing. In, 2000-1-1Elsevier), p. 40–46.

45. Charlton, M. (2009). Obesity, hyperlipidemia, and metabolic syndrome. Liver Transplant. 15, S83–S89.

46. Ferrero, R., Rainer, P.Y., Rumpler, M., Russeil, J., Zachara, M., Pezoldt, J., van Mierlo, G., Gardeux, V., Saelens, W., and Alpern, D. (2024). A human omentum-specific mesothelial-like stromal population inhibits adipogenesis through IGFBP2 secretion. Cell Metab.

47. Fullerton, M.D., Galic, S., Marcinko, K., Sikkema, S., Pulinilkunnil, T., Chen, Z., O’Neill, H.M., Ford, R.J., Palanivel, R., and O’Brien, M. (2013). Single phosphorylation sites in Acc1 and Acc2 regulate lipid homeostasis and the insulin-sensitizing effects of metformin. Nat. Med. 19, 1649–1654.

48. Gross, B., Pawlak, M., Lefebvre, P., and Staels, B. (2017). PPARs in obesity-induced T2DM, dyslipidaemia and NAFLD. Nat. Rev. Endocrinol. 13, 36–49.

49. Ju, U., Jeong, D., Seo, J., Park, J.B., Park, J., Suh, K., Kim, J.B., and Chun, Y. (2020). Neddylation of sterol regulatory element-binding protein 1c is a potential therapeutic target for nonalcoholic fatty liver treatment. Cell Death Dis. 11, 283.

50. Liu, B., Liu, X., Liang, Z., and Wang, J. (2021). Gut microbiota in obesity. World J. Gastroenterol. 27, 3837.

51. Takagi, T., Naito, Y., Kashiwagi, S., Uchiyama, K., Mizushima, K., Kamada, K., Ishikawa, T., Inoue, R., Okuda, K., and Tsujimoto, Y. (2020). Changes in the gut microbiota are associated with hypertension, hyperlipidemia, and type 2 diabetes mellitus in Japanese subjects. Nutrients 12, 2996.

52. Gupta, V.K., Kim, M., Bakshi, U., Cunningham, K.Y., Davis Iii, J.M., Lazaridis, K.N., Nelson, H., Chia, N., and Sung, J. (2020). A predictive index for health status using species-level gut microbiome profiling. Nat. Commun. 11, 4635.

53. Bourdeau-Julien, I., Castonguay-Paradis, S., Rochefort, G., Perron, J., Lamarche, B., Flamand, N., Di Marzo, V., Veilleux, A., and Raymond, F. (2023). The diet rapidly and differentially affects the gut microbiota and host lipid mediators in a healthy population. Microbiome 11, 26.

54. Dunham, I. (2018). Human genes: Time to follow the roads less traveled? PLoS Biol. 16, e3000034.

55. Kustatscher, G., Collins, T., Gingras, A., Guo, T., Hermjakob, H., Ideker, T., Lilley, K.S., Lundberg, E., Marcotte, E.M., and Ralser, M. (2022). Understudied proteins: opportunities and challenges for functional proteomics. Nat. Methods 19, 774–779.

56. Unsal, S., Atas, H., Albayrak, M., Turhan, K., Acar, A.C., and Doğan, T. (2022). Learning functional properties of proteins with language models. Nat. Mach. Intell. 4, 227–245.

57. Wan, F., Torres, M.D., Peng, J., and de la Fuente-Nunez, C. (2024). Deep-learning-enabled antibiotic discovery through molecular de-extinction. Nat. Biomed. Eng., 1–18.

58. Santos-Júnior, C.D., Torres, M.D., Duan, Y., Del Río, Á.R., Schmidt, T.S., Chong, H., Fullam, A., Kuhn, M., Zhu, C., and Houseman, A. (2024). Discovery of antimicrobial peptides in the global microbiome with machine learning. Cell.

59. Melo, M.C., Maasch, J.R., and de la Fuente-Nunez, C. (2021). Accelerating antibiotic discovery through artificial intelligence. Commun. Biol. 4, 1050.

60. Adessi, C., and Soto, C. (2002). Converting a peptide into a drug: strategies to improve stability and bioavailability. Curr. Med. Chem. 9, 963–978.

61. Kyte, J. (2006). Structure in protein chemistry (Garland Science).

62. Sewald, N., and Jakubke, H. (2015). Peptides: chemistry and biology (John Wiley & Sons).

63. Das, P., Sercu, T., Wadhawan, K., Padhi, I., Gehrmann, S., Cipcigan, F., Chenthamarakshan, V., Strobelt, H., Dos Santos, C., and Chen, P. (2021). Accelerated antimicrobial discovery via deep generative models and molecular dynamics simulations. Nat. Biomed. Eng. 5, 613–623.

64. Teufel, F., Gíslason, M.H., Almagro Armenteros, J.J., Johansen, A.R., Winther, O., and Nielsen, H. (2023). GraphPart: homology partitioning for biological sequence analysis. NAR Genom. Bioinform. 5, lqad088.

65. Rives, A., Meier, J., Sercu, T., Goyal, S., Lin, Z., Liu, J., Guo, D., Ott, M., Zitnick, C.L., and Ma, J. (2021). Biological structure and function emerge from scaling unsupervised learning to 250 million protein sequences. Proceedings of the National Academy of Sciences 118, e2016239118.

66. Høie, M.H., Kiehl, E.N., Petersen, B., Nielsen, M., Winther, O., Nielsen, H., Hallgren, J., and Marcatili, P. (2022). NetSurfP-3.0: accurate and fast prediction of protein structural features by protein language models and deep learning. Nucleic. Acids. Res. 50, W510–W515.

67. Wishart, D.S., Feunang, Y.D., Guo, A.C., Lo, E.J., Marcu, A., Grant, J.R., Sajed, T., Johnson, D., Li, C., and Sayeeda, Z. (2018). DrugBank 5.0: a major update to the DrugBank database for 2018. Nucleic. Acids. Res. 46, D1074–D1082.

68. Kim, S., Chen, J., Cheng, T., Gindulyte, A., He, J., He, S., Li, Q., Shoemaker, B.A., Thiessen, P.A., and Yu, B. (2021). PubChem in 2021: new data content and improved web interfaces. Nucleic. Acids. Res. 49, D1388–D1395.

69. Carbon, S., Ireland, A., Mungall, C.J., Shu, S., Marshall, B., Lewis, S., Amigo, H., and Web, P.W.G. (2009). AmiGO: online access to ontology and annotation data. Bioinformatics 25, 288–289.

70. Sherman, B.T., Hao, M., Qiu, J., Jiao, X., Baseler, M.W., Lane, H.C., Imamichi, T., and Chang, W. (2022). DAVID: a web server for functional enrichment analysis and functional annotation of gene lists (2021 update). Nucleic. Acids. Res. 50, W216–W221.

71. Mészáros, B., Erdős, G., and Dosztányi, Z. (2018). IUPred2a: context-dependent prediction of protein disorder as a function of redox state and protein binding. Nucleic. Acids. Res. 46, W329–W337.

72. Rice, P., Longden, I., and Bleasby, A. (2000). EMBOSS: the European molecular biology open software suite. Trends Genet. 16, 276–277.

73. Moriwaki, H., Tian, Y., Kawashita, N., and Takagi, T. (2018). Mordred: a molecular descriptor calculator. J. Cheminformatics 10, 1–14.

74. Wei, L., Ye, X., Sakurai, T., Mu, Z., and Wei, L. (2022). ToxIBTL: prediction of peptide toxicity based on information bottleneck and transfer learning. Bioinformatics 38, 1514–1524.

75. Li, Z., Zhang, B., Wang, N., Zuo, Z., Wei, H., and Zhao, F. (2023). A novel peptide protects against diet-induced obesity by suppressing appetite and modulating the gut microbiota. Gut 72, 686–698.

