## Supplementary Figure 1-8 (termed as Figure S1-S8) for "Discovery of potent oligopeptides for various metabolic diseases using deep learning"

**Supplementary figures**

**Figure S1.** The deep learning model architecture.

**Figure S2.** The determination of five indications and their molecular function fingerprints.

**Figure S3.** Distribution of identified oligopeptides and their overlapping with known peptides across five indications.

**Figure S4.** Scratch, CCK-8 and tube formation assays of HUVEC cells treated with oligopeptides.

**Figure S5.** Oil Red O staining of adipogenic-induced 3T3-L1 cells treated with oligopeptides.

**Figure S6.** *In vivo* pharmacodynamic and safety evaluation of TP6 in comparison with first-line drugs for obesity and hyperlipidemia.

**Figure S7.** Evaluation of the effects of selected peptides across five indications, compared to randomly generated peptides and scrambled controls.

**Figure S8.** Results of qPCR analysis for the five indications.

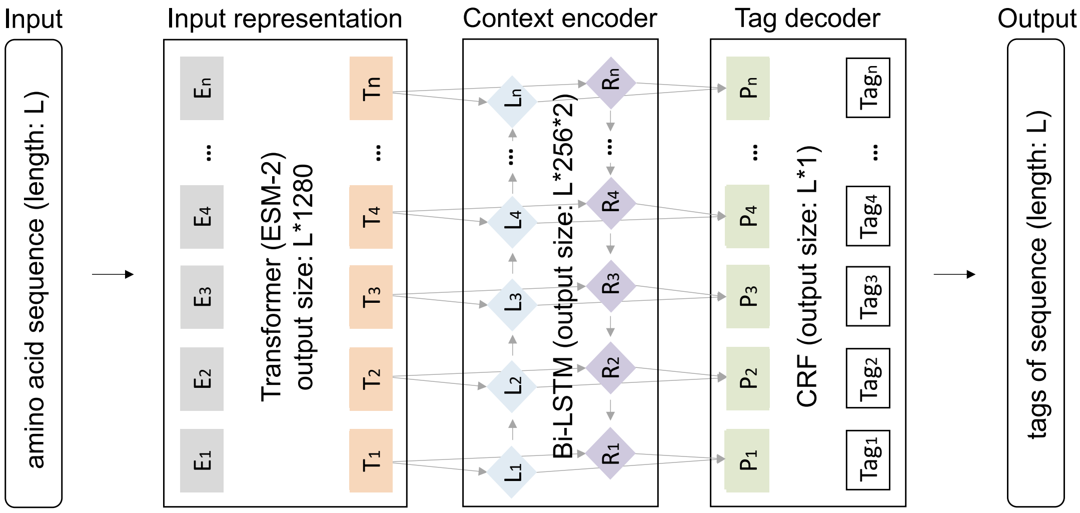


**Figure S1. The deep learning model architecture and data-handling process.** The deep learning model architecture. The deep learning model is designed to identify biopeptide candidates from input amino acid (AA) sequences. It comprises three main components: 1) Input Representation: The pre-trained Transformer-based large language model, ESM-2, is fine-tuned to incorporate the global evolutionary information and general semantic patterns of proteins. 2) Context Encoder: Bidirectional Long Short-Term Memory (Bi-LSTM) captures context dependencies for inferring the boundary between interested subsequence and flanking sequences. 3) Tag Decoder: A Conditional Random Field (CRF) assigns a tag to each amino acid, which is subsequently used to extract the predicted oligopeptide from the protein. For the *i*-th AA, the process involves: a) $E_i$, the input embedding; b) $T_i$, the encoded output from the Transformer model; c) $L_i$, incorporating information of left context and itself; d) $R_i$, incorporating information of right context and itself; e) $P_i$, the probability vector representing the four tags ('B', 'I', 'E', and 'O' indicating the beginning, inner, end, and outside of biopeptides, respectively).

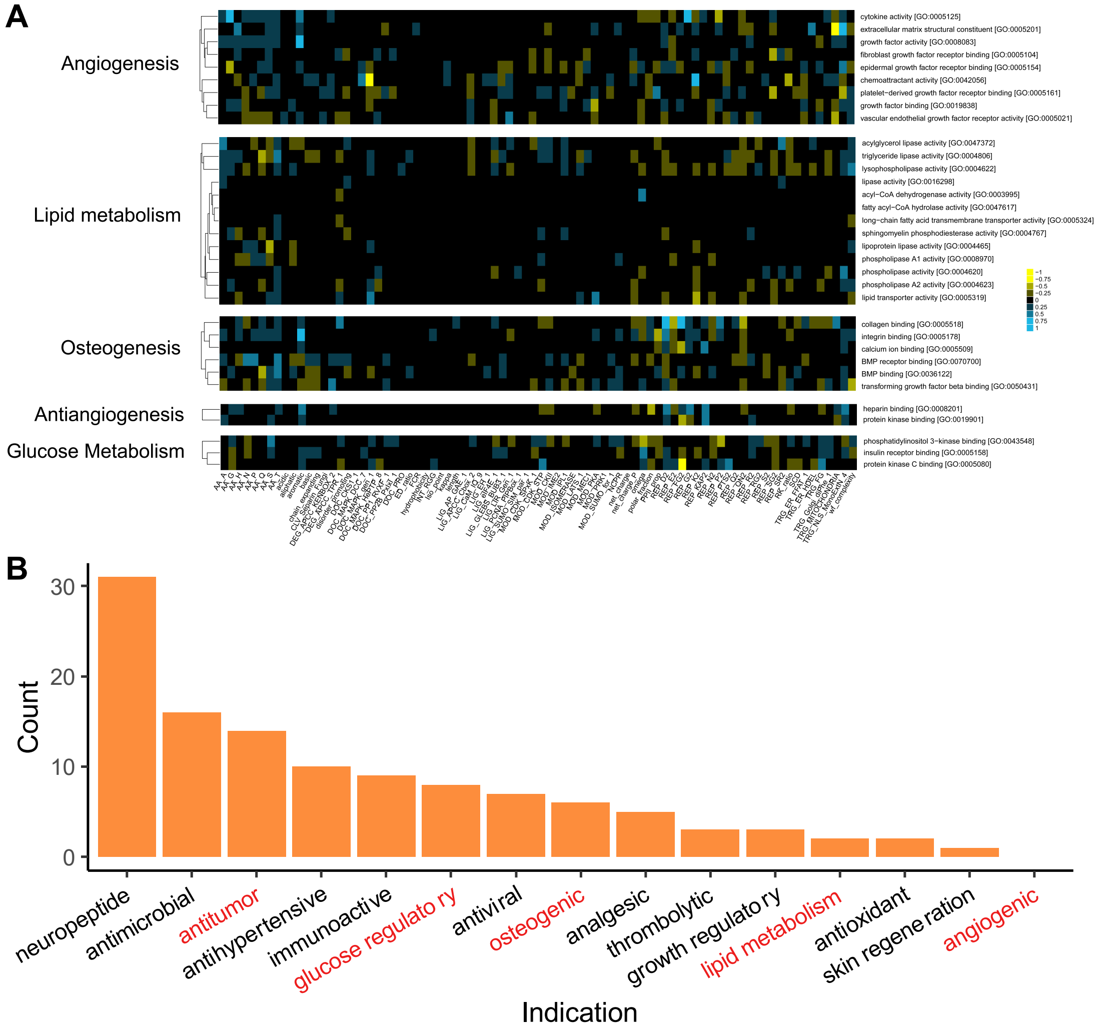


**Figure S2. The determination of five indications and their molecular function fingerprints. (A)** Molecular function fingerprints (MFFs) for the five indications. Molecular functions of the five indications (y-axis) are plotted against molecular features (x-axis) as determined by FAIDR. The colors represent the extent to which the molecular feature is associated with a certain molecular function. Specifically, blue indicates a positive correlation, and purple indicates a negative correlation. **(B)** The distribution of marketed peptide drugs by indication. Considering broad relevance and potential clinical utility, we selected five indications for therapeutic oligopeptide lead discovery: angiogenesis, lipid metabolism, osteogenesis, glucose metabolism, and anti-angiogenesis (red letters).

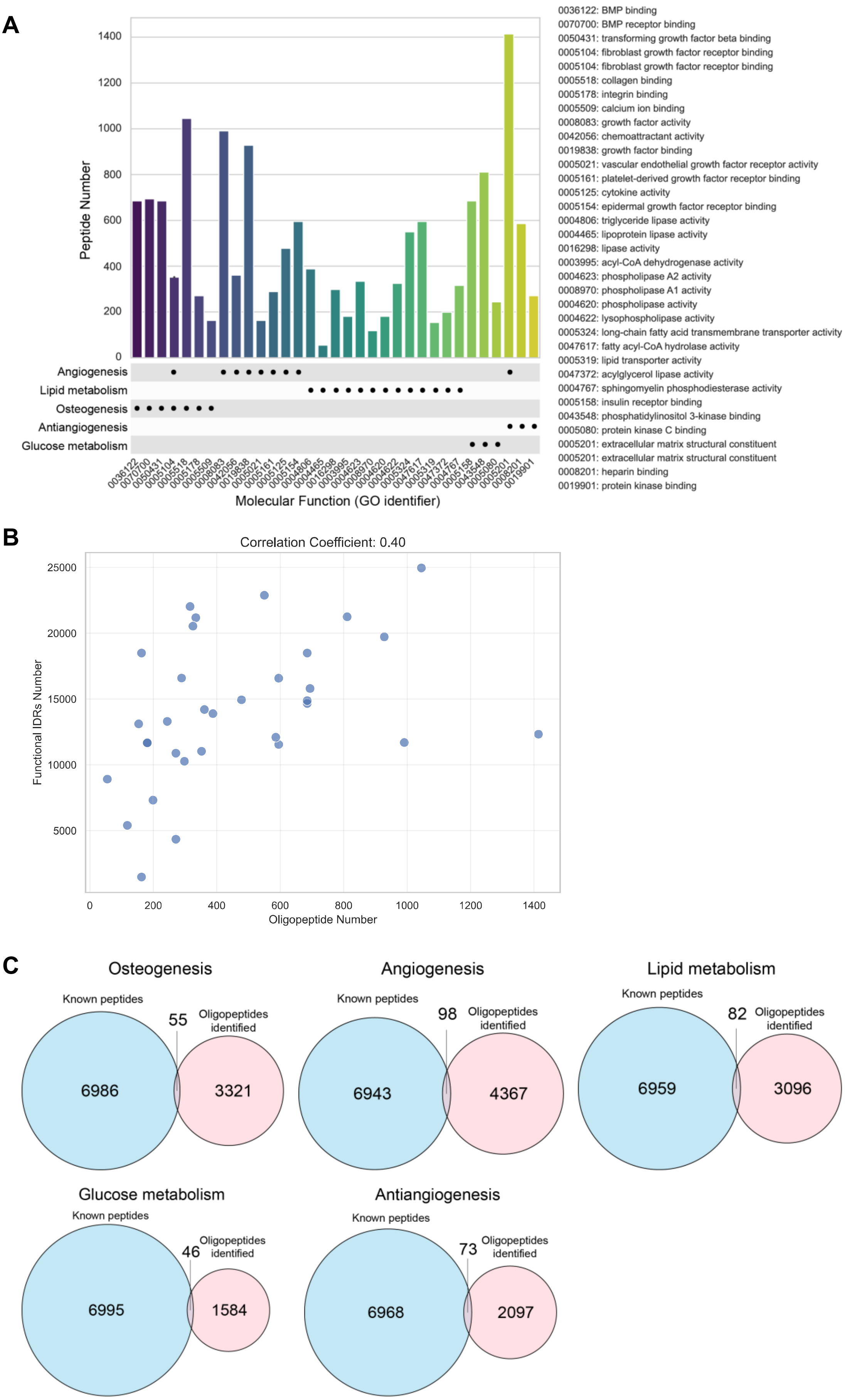
A
Peptide Number
Angiogenesis
Lipid metabolism
Osteogenesis
Antiangiogenesis
Glucose metabolism
Molecular Function (GO identifier)
0036122: BMP binding
0070700: BMP receptor binding
0050431: transforming growth factor beta binding
0005104: fibroblast growth factor receptor binding
0005104: fibroblast growth factor receptor binding
0005518: collagen binding
0005178: integrin binding
0005509: calcium ion binding
0008083: growth factor activity
0042056: chemoattractant activity
0019838: growth factor binding
0005021: vascular endothelial growth factor receptor activity
0005161: platelet-derived growth factor receptor binding
0005125: cytokine activity
0005154: epidermal growth factor receptor binding
0004806: triglyceride lipase activity
0004465: lipoprotein lipase activity
0016298: lipase activity
0003995: acyl-CoA dehydrogenase activity
0004623: phospholipase A2 activity
0008970: phospholipase A1 activity
0004620: phospholipase activity
0004622: lysophospholipase activity
0005324: long-chain fatty acid transmembrane transporter activity
0047617: fatty acyl-CoA hydrolase activity
0005319: lipid transporter activity
0047372: acylglycerol lipase activity
0004767: sphingomyelin phosphodiesterase activity
0005158: insulin receptor binding
0043548: phosphatidylinositol 3-kinase binding
0005080: protein kinase C binding
0005201: extracellular matrix structural constituent
0005201: extracellular matrix structural constituent
0008201: heparin binding
0019901: protein kinase binding
B
Correlation Coefficient: 0.40
Functional IDRs Number
Oligopeptide Number
C
Osteogenesis
Known peptides
55
Oligopeptides identified
6986
3321
Angiogenesis
Known peptides
98
Oligopeptides identified
6943
4367
Lipid metabolism
Known peptides
82
Oligopeptides identified
6959
3096
Glucose metabolism
Known peptides
46
Oligopeptides identified
6995
1584
Antiangiogenesis
Known peptides
73
Oligopeptides identified
6968
2097

**Figure S3. Distribution of identified oligopeptides and their overlapping with known peptides across five indications. (A)** Distribution of all 33 molecular functions across five indications for identified oligopeptides. Therapeutic oligopeptide candidates show significant difference in distribution between different molecular functions, likely influenced by the number of functional IDRs belonging to the corresponding molecular functions (show in panel **B**). **(B)** Number distribution and correlation analysis of functional IDRs and oligopeptides belonging to these molecular functions. The correlation coefficient is 0.4. **(C)** Overlap of identified oligopeptides with known peptides in the training dataset for each disease indication.

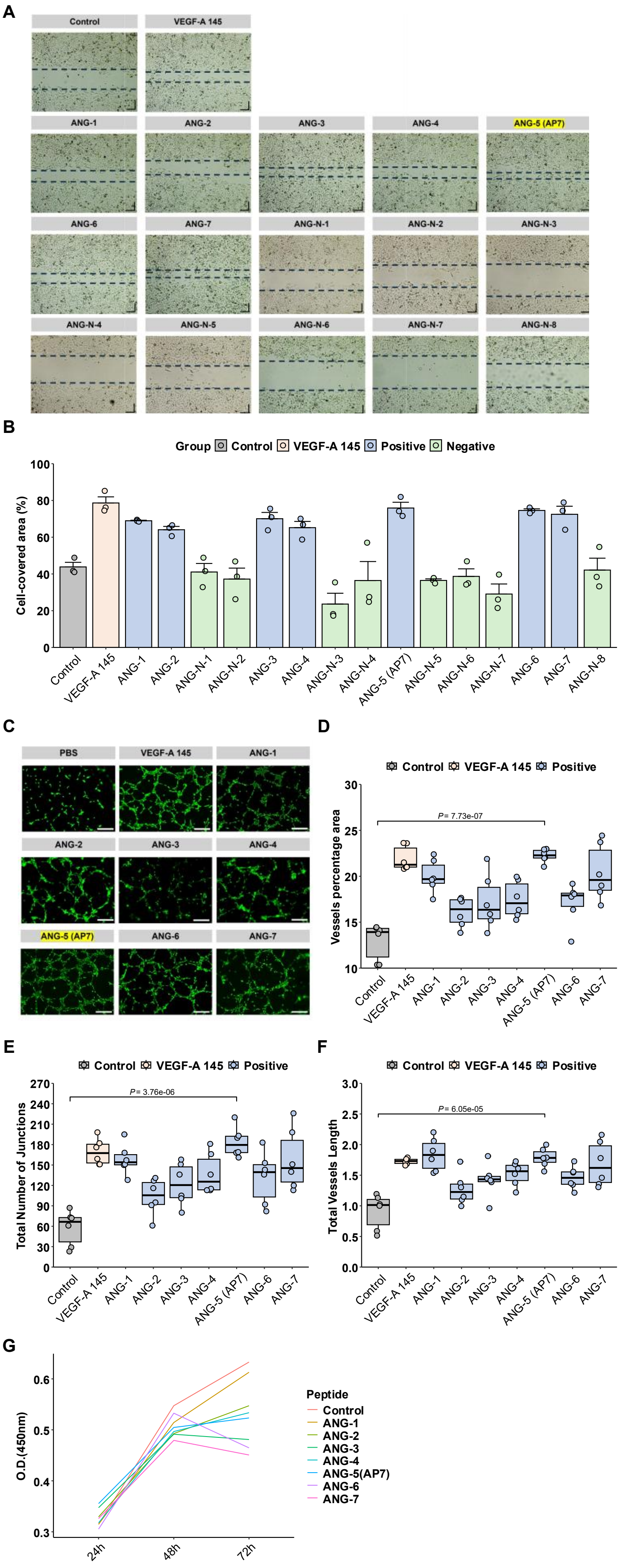
A
Control
VEGF-A 145
ANG-1
ANG-2
ANG-3
ANG-4
ANG-5 (AP7)
ANG-6
ANG-7
ANG-N-1
ANG-N-2
ANG-N-3
ANG-N-4
ANG-N-5
ANG-N-6
ANG-N-7
ANG-N-8
B
Group Control VEGF-A 145 Positive Negative
Cell-covered area (%)
C
PBS
VEGF-A 145
ANG-1
ANG-2
ANG-3
ANG-4
ANG-5 (AP7)
ANG-6
ANG-7
D
Control VEGF-A 145 Positive
Vessels percentage area
P= 7.73e-07
E
Control VEGF-A 145 Positive
Total Number of Junctions
P= 3.76e-06
F
Control VEGF-A 145 Positive
Total Vessels Length
P= 6.05e-05
G
O.D.(450nm)
24h
48h
72h
Peptide
Control
ANG-1
ANG-2
ANG-3
ANG-4
ANG-5(AP7)
ANG-6
ANG-7

**Figure S4. Scratch, CCK-8 and tube formation assays of HUVEC cells treated with oligopeptides. (A)** Quantified cell coverage of HUVEC cells is measured using ImageJ after treatment with 0.1 ug·mL$^{-1}$ oligopeptides. **(B)** Representative images of scratch assays of HUVEC cells treated with 0.1 ug·mL$^{-1}$ oligopeptides**. (C-F**) Tube formation assays of HUVEC cells treated with 0.2 ug·mL$^{-1}$ oligopeptides. **(G)** CCK-8 assay of HUVEC cells treated with 0.1 ug·mL$^{-1}$ oligopeptides. The yellow-highlighted candidate, namely ANG-5, is AP7 selected for subsequent *in vivo* validations. The experiment was repeated three times independently. Scale bar = 10μm. *$P < 0.05$, **$P < 0.01$, ***$P < 0.001$, ****$P < 0.0001$. Significant differences between two groups (wild type and each oligopeptide treatment) are determined by one-way ANOVA. Data are presented as the means ± SEM. Source data are provided as a Source Data file.

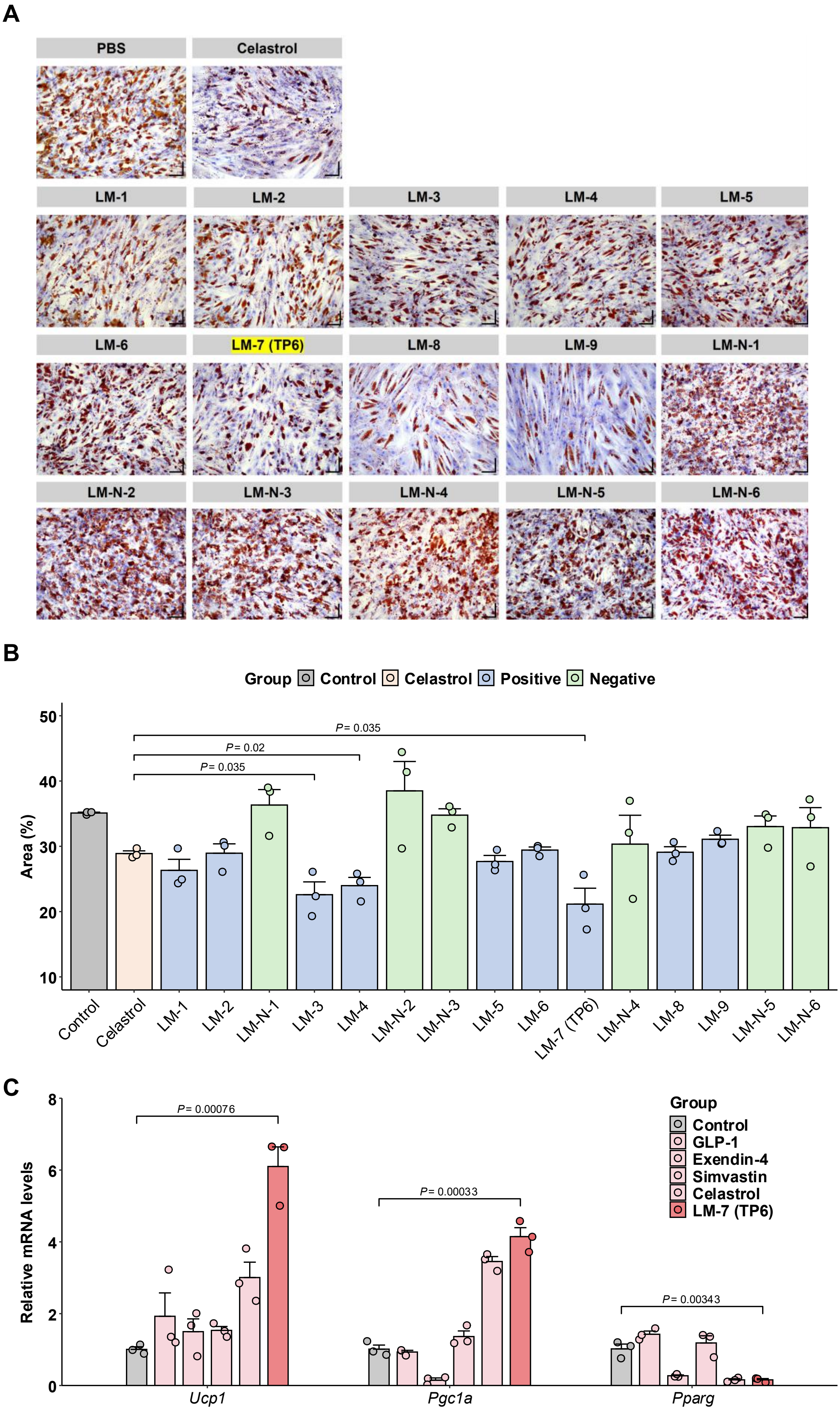
A
PBS
Celastrol
LM-1
LM-2
LM-3
LM-4
LM-5
LM-6
LM-7 (TP6)
LM-8
LM-9
LM-N-1
LM-N-2
LM-N-3
LM-N-4
LM-N-5
LM-N-6
B
Group Control Celastrol Positive Negative
Area (%)
50
40
30
20
10
P = 0.035
P = 0.02
P = 0.035
Control
Celastrol
LM-1
LM-2
LM-N-1
LM-3
LM-4
LM-N-2
LM-N-3
LM-5
LM-6
LM-7 (TP6)
LM-N-4
LM-8
LM-9
LM-N-5
LM-N-6
C
Relative mRNA levels
8
6
4
2
0
P = 0.00076
P = 0.00033
P = 0.00343
Ucp1
Pgc1a
Pparg
Group
Control
GLP-1
Exendin-4
Simvastin
Celastrol
LM-7 (TP6)

**Figure S5. Oil Red O staining of adipogenic-induced 3T3-L1 cells treated with oligopeptides. (A)** Quantified area of Oil red O-stained adipogenic-induced 3T3-L1 cells treated with 0.1 $ug{\cdot}mL^{-1}$ oligopeptides is measured using ImageJ. **(B)** Oil red O-stained adipogenic-induced 3T3-L1 cells**.** The yellow highlighted candidate, namely LM-7, is TP6, selected for subsequent *in vivo* validations. **(C)** qPCR analysis of *Ucp1*, *Pgc1a*, and *Pparγ* gene expression in cells stimulated with LM-7 (TP6) and four positive controls (GLP-1, Exendin-4, Simvastatin, and Celastrol). The experiment is repeated three times independently. Scale bar = 25μm. Significance markers: $^*P < 0.05$, $^{**}P < 0.01$; ns: not significant. Significant differences between two groups (wild type and each oligopeptides treated) are determined by one-way ANOVA. Data are presented as the means ± SEM. Source data is provided as a Source Data file.

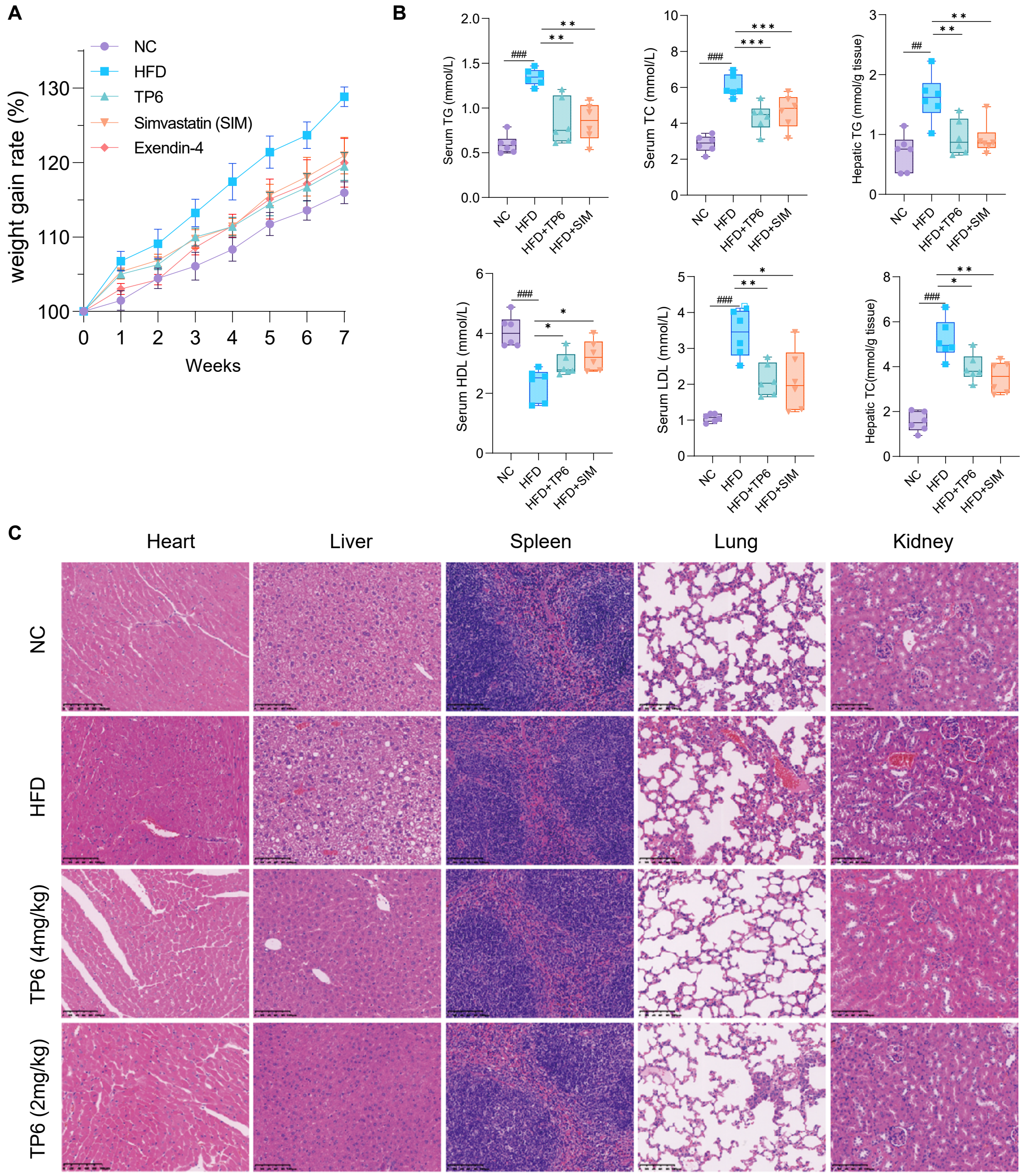


**Figure S6. *In vivo* pharmacodynamic and safety evaluation of TP6 in comparison with first-line drugs for obesity and hyperlipidemia. (A)** Grams of weight gain rate measured over time. **(B)** Serum TG, TC, HDL and HDL contents,and hepatic TG and TC contents. 30 C57/Bl6J mice were divided into five groups (n=6/group): NC, HFD, HFD+TP6 (4 mg/kg), HFD+Exendin-4 (10 μg/kg), HFD+Simvastatin (1.5mg/kg). **(C)** Representative H&E-stained images of heart, liver, spleen, lung, and kidney tissues collected from mice at the completion of the experiment. Scale bar: 100 μm. The experiment lasted for 7 weeks. HFD, high fat diet. SIM, Simvastatin. $P < 0.05$ (* or #); $P < 0.01$ (** or ##); $P < 0.001$ (*** or ###).

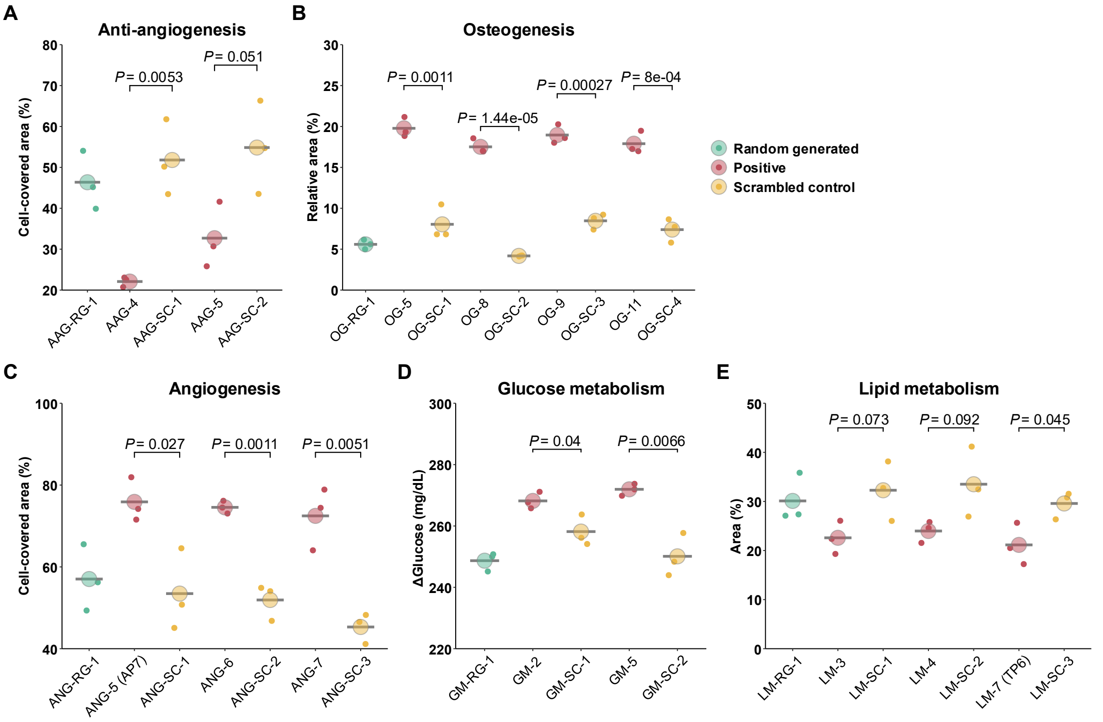


**Figure S7. Evaluation of the effects of selected peptides across five indications, compared to randomly generated peptides and scrambled controls**. Red dots represent positive peptides identified through Deepeptide, green dots indicate randomly generated peptides, and yellow dots correspond to scrambled controls. Each data point represents an individual biological replicate. The five indications assessed are anti-angiogenesis (**A**), Osteogenesis (**B**), Angiogenesis (**C**), Glucose metabolism (**D**), and Lipid metabolism (**E**).

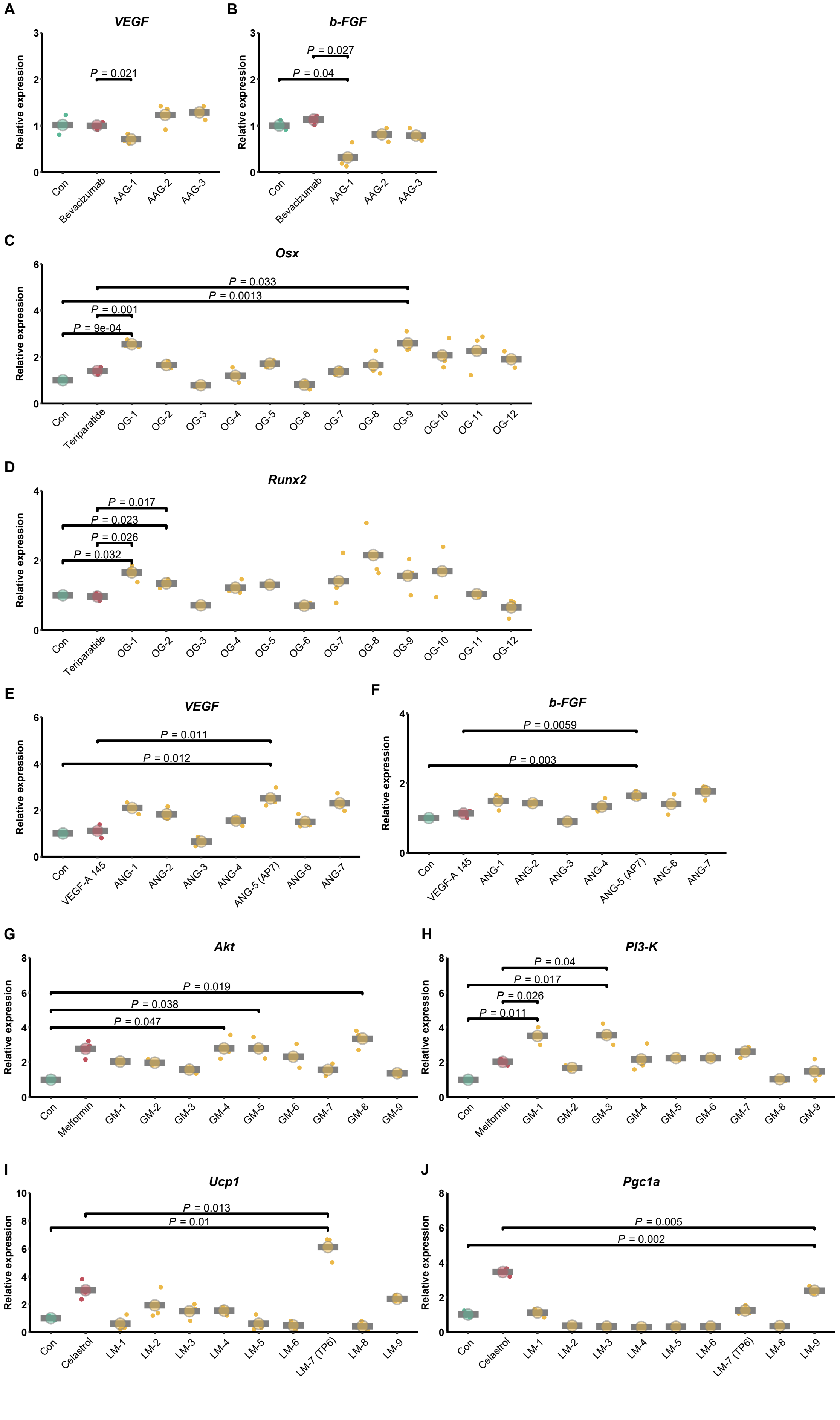
A
VEGF
Relative expression
P = 0.021
Con
Bevacizumab
AAG-1
AAG-2
AAG-3
B
b-FGF
Relative expression
P = 0.027
P = 0.04
Con
Bevacizumab
AAG-1
AAG-2
AAG-3
C
Osx
Relative expression
P = 0.033
P = 0.0013
P = 0.001
P = 9e-04
Con
Teriparatide
OG-1
OG-2
OG-3
OG-4
OG-5
OG-6
OG-7
OG-8
OG-9
OG-10
OG-11
OG-12
D
Runx2
Relative expression
P = 0.017
P = 0.023
P = 0.026
P = 0.032
Con
Teriparatide
OG-1
OG-2
OG-3
OG-4
OG-5
OG-6
OG-7
OG-8
OG-9
OG-10
OG-11
OG-12
E
VEGF
Relative expression
P = 0.011
P = 0.012
Con
VEGF-A 145
ANG-1
ANG-2
ANG-3
ANG-4
ANG-5 (AP7)
ANG-6
ANG-7
F
b-FGF
Relative expression
P = 0.0059
P = 0.003
Con
VEGF-A 145
ANG-1
ANG-2
ANG-3
ANG-4
ANG-5 (AP7)
ANG-6
ANG-7
G
Akt
Relative expression
P = 0.019
P = 0.038
P = 0.047
Con
Metformin
GM-1
GM-2
GM-3
GM-4
GM-5
GM-6
GM-7
GM-8
GM-9
H
PI3-K
Relative expression
P = 0.04
P = 0.017
P = 0.026
P = 0.011
Con
Metformin
GM-1
GM-2
GM-3
GM-4
GM-5
GM-6
GM-7
GM-8
GM-9
I
Ucp1
Relative expression
P = 0.013
P = 0.01
Con
Celastrol
LM-1
LM-2
LM-3
LM-4
LM-5
LM-6
LM-7 (TP6)
LM-8
LM-9
J
Pgc1a
Relative expression
P = 0.005
P = 0.002
Con
Celastrol
LM-1
LM-2
LM-3
LM-4
LM-5
LM-6
LM-7 (TP6)
LM-8
LM-9

**Figure S8. Results of qPCR analysis for the five indications.** qPCR analysis was performed on all 40 positive oligopeptides across the following indications: antiangiogenesis (**A-B**), osteogenesis (**C-D**), angiogenesis (**E-F**), lipid metabolism (**G-H**), and glucose metabolism (**I-J**). As shown, the majority of identified positive oligopeptides promote osteogenesis by regulating *Osx* or *Runx2*, enhance angiogenesis via modulation of *b-FGF* or *VEGF*, modulate lipid metabolism through *Pgc1a* or *Ucp1*, regulate glucose metabolism via *Akt* or *PI3-K*, and inhibit angiogenesis by affecting *b-FGF* or *VEGF*.
